# Loss of PIKfyve drives the spongiform degeneration in prion diseases

**DOI:** 10.1101/2021.01.08.425896

**Authors:** Asvin KK Lakkaraju, Karl Frontzek, Emina Lemes, Uli Herrmann, Marco Losa, Rajlakshmi Marpakwar, Adriano Aguzzi

## Abstract

Brain-matter vacuolation is a defining trait of all prion diseases, yet its cause is unknown. Here we report that prion infection and prion-mimetic antibodies deplete the phosphatidylinositol kinase PIKfyve in mouse brains, cultured cells, organotypic brain slices, and in brains of Creutzfeldt-Jakob Disease victims. We found that PIKfyve, an inositol kinase involved endolysosomal maturation, is acylated by zDHHC9 and zDHHC21, whose juxtavesicular topology is disturbed by prion infection, resulting in PIKfyve deacylation and destabilization. A protracted unfolded protein response (UPR), typical of prion diseases, also induced PIKfyve deacylation and degradation. Conversely, UPR antagonists restored PIKfyve levels in prion-infected cells. Overexpression of zDHHC9 and zDHHC21, administration of the antiprion polythiophene LIN5044, or supplementation with the PIKfyve reaction product PI(3,5)P_2_, suppressed prion-induced vacuolation. Thus, PIKfyve emerges as a central mediator of vacuolation and neurotoxicity in prion diseases.

---

In prion diseases, intraneuronal vacuoles gradually give rise to spongiosis, an extensive sponge-like state that eventually occupies much of the cerebral cortex. These vacuoles may arise from lysosomes, whose function can be impaired by prion infections^1,2^. Lysosome maturation is controlled by the phosphoinositide diphosphate PI(3,5)P_2_ generated by the PIKfyve kinase. PIKfyve forms a complex with the phosphatase FIG4 and the scaffolding protein VAC14 on the cytosolic face of endosomes. Ablation of any of these proteins alters the PI(3,5)P_2_ levels, resulting in enlarged endolysosomes reminiscent of prion-induced spongiosis^3-5^. We therefore asked whether PIKfyve is involved in prion-induced spongiosis.

Six C57BL/6 wild-type mice were inoculated intracerebrally with RML scrapie prions (Rocky Mountain Laboratory strain, passage 6) and sacrificed at terminal disease 184 ± 16 days post-infection (dpi). For control, we inoculated 4 mice with non-infectious brain homogenate (NBH, 190 dpi). Prion-infected brains showed a profound reduction of PIKfyve, but not of FIG4 and VAC14 (Fig. 1A). PIKfyve was not affected in other organs (Fig. S1A). PIKfyve levels were also reduced in brains of mice infected with the ME7 strain of prions (Fig. 1B) and in RML-infected PrP^C^-overexpressing *tg*a*20* mice^6^ (Fig. S1B). Tumor susceptibility gene 101 (Tsg101), superoxide dismutase-2 and Mahogunin-1, which have been implicated in vacuolating brain diseases including genetic spongiform encephalopathies^7-9^, were unchanged (Fig. S1C). PIKfyve levels were decreased at 120 dpi and was profoundly depleted at the terminal stage (Fig. 1C-D). PIKfyve is highly expressed in neurons^10^, yet the neuronal proteins NeuN and synaptophysin were unaffected (Fig. 1C), suggesting that its loss did not reflect the altered brain tissue composition of scrapie-sick mice.

**Figure 1:**
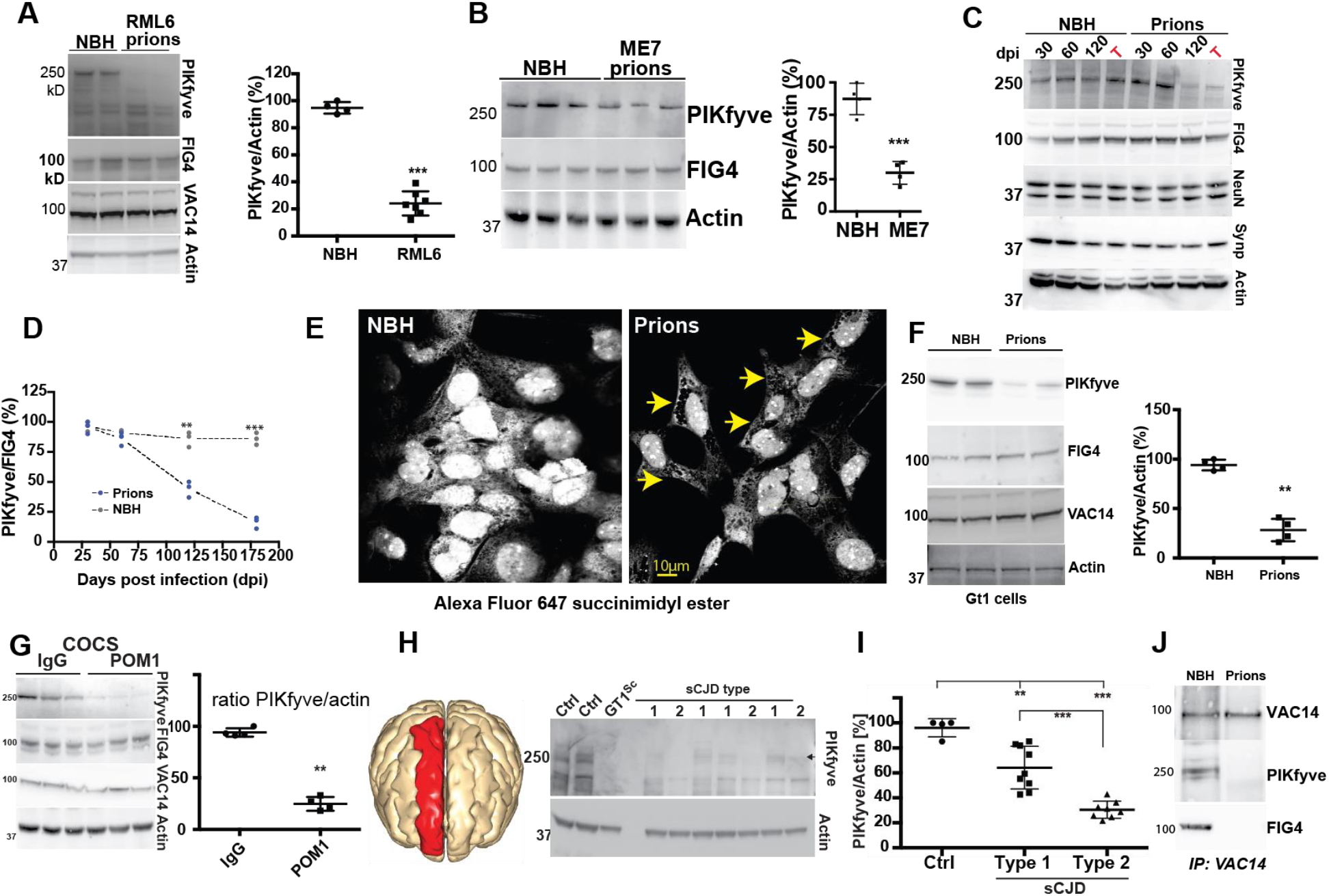
PIKfyve is universally depleted in prion diseases. **A**: Conspicuous downregulation of PIKfyve, but not FIG4 or VAC14, in brain lysates of terminally sick mice intracerebrally inoculated with prions (RML). Control: mice inoculated with non-infectious brain homogenate (NBH). Right: Quantification of immunoblot. Each dot represents one mouse. *: p0.05; **: p<0.01; ***: p<0.001 here and henceforth; unpaired t-test. **B**: PIKfyve is selectively downregulated in mice infected with ME7 prions. Right: Quantification. Each dot represents one individual (n=4/group). **C:** Time course of PIKfyve downregulation in brains of prion-infected mice (3 mice/time point). PIKfyve (but not FIG4, NeuN and synaptophysin) was suppressed between 60 and 120 dpi. T: terminal disease stage. **D**: The PIKfyve/FIG4 ratio gradually decreased in prion-infected mice (unpaired t-test). **E**: Prion-infected Gt1 cells (75 dpi) stained with Alexa-Fluor647-succinimidyl ester which stains vacuoles negatively (arrows). **F:** Loss of PIKfyve, but not FIG4 and VAC14, in prion-infected Gt1 cells. Right: Quantification of immunoblots. Each dot represents an independent experiment. **G:** *tg*a*20* COCS were treated with pooled IgG or POM1 (72 hours). POM1 suppressed PIKfyve but not FIG4 and VAC14. Right: Quantification of immunoblots. Each dot represents an independent experiment. **H:** Immunoblots of human CJD brains (red: sampled region). Gt1^Sc^: prion-infected Gt1 cells. **I**: Quantification of immunoblots revealed moderate and substantial PIKfyve loss in Type-1 and Type-2 CJD, respectively (unpaired t-test). **J:** Mouse brain homogenates were immunoprecipitated with anti-VAC14 antibodies. FIG4 and PIKfyve did not co-precipitate in prion-infected brains.

We then infected murine Gt1 cells^11^ with RML prions. At 75 dpi we observed the appearance of cytoplasmic vacuoles accompanied by a substantial reduction of PIKfyve, but not of FIG4 or VAC14 (Fig. 1E-F). PrP depletion from prion-infected Gt1 cells by RNAi at 70 dpi largely suppressed cytoplasmic vacuolation (Fig. S1D). Next, we administered RML prions to cerebellar organotypic cultured slices (COCS) from 9-day old *tg*a*20* mice^12^. At 45 dpi we observed neurodegeneration accompanied by conspicuous PIKfyve depletion (Fig. S1E-F). Finally, we exposed COCS for 72 hours to the antibody POM1 which induces PrP^C^-dependent neurotoxicity^13^ similarly to prion infections^14^. Again, this treatment depleted PIKfyve from COCS (Fig. 1G). We then asked if PIKfyve is lost in human sporadic Creutzfeldt-Jakob disease (sCJD). Differential proteolysis of the misfolded prion protein (PrP^Sc^) discriminates two types of sCJD with distinct clinical courses and histopathological characteristics^15^. We investigated 9 and 8 frontal cortex samples (Fig. S1G-H) from patients who succumbed from type-1 and type-2 sCJD, respectively, as well as 4 non-CJD controls, collected between 2002 and 2010 (Table S1). PIKfyve levels were reduced in all cases, but the reduction was more profound in type-2 sCJD samples (Fig. 1H-I) which also showed more extensive vacuolation (Fig. S1H). Along with the earlier onset of disease in Type-2 CJD (average: 57 vs. 63 years)^16^, this suggests that PIKfyve depletion may be a determinant of toxicity. Therefore, PIKfyve depletion is a feature of all investigated instances of prion infections.

We then studied the consequences of prion infection onto the composition of the PIKfyve tripartite complex. While VAC14 and FIG4 levels were not affected by prion infection, immunoprecipitation of VAC14 captured both FIG4 and PIKfyve from NBH-inoculated brains, but not from prion-infected brains (Fig. 1J). Conversely, the siRNA knockdown of FIG4 or VAC14 in Gt1 cells using RNAi caused a significant PIKfyve reduction. The knockdown of FIG4 reduced VAC14 levels but the VAC14 knockdown did not alter FIG4(Fig. S1I). Hence the prion-induced PIKfyve depletion destabilizes the association between FIG4 and VAC14 without reducing their concentration.

Next, we investigated the mechanism of PIKfyve suppression by prions. Prion-infected mice (Fig. S1J), as well as prion-infected or POM1-treated COCS (Figs. S1K and S1L, respectively) showed no alterations in PIKfyve, FIG4 and VAC14 brain mRNA levels. The relative expression of PIKfyve mRNA splice variants was similar between RML-infected and control mouse brains (Fig. S1M-N), and exploration of a splice alteration database during the progression of prion disease^17^ did not identify any differential expression of PIKfyve isoforms. Hence PIKfyve suppression occurs post-transcriptionally.

Prion diseases induce a chronic unfolded-protein response (UPR) in the endoplasmic reticulum (ER) leading to eIF2α phosphorylation and translational suppression^18^, and UPR inhibitors can mitigate the spongiosis^19^. We first compared the evolution of the UPR and of PIKfyve depletion in various prion models. In prion-infected mouse brains, eIF2α phosphorylation was detectable at 30 dpi, whereas PIKfyve decreased gradually starting at 60 dpi (Fig. 2A-B). In COCS, eIF2α became phosphorylated 8h after exposure to POM1, yet PIKfyve was depleted only after 72 hours (Fig. 2C, S2A). Hence a chronic UPR precedes downregulation of PIKfyve in vivo and ex vivo. We then induced an acute UPR in Gt1 cells with thapsigargin (0.5 µM) resulting in depletion of PIKfyve substantially but transiently (Fig. 2D, S2B-C). We then assessed the half-life of PIKfyve in thapsigargin or DMSO-treated Gt1 cells (4 hours) using a [^35^S]methionine/cysteine pulse (20 min) followed by chasing in ^35^S-free culture medium. Thapsigargin reduced the half-life of immunopurified [^35^S]PIKfyve from 79 to 38 min (Fig. 2E-F).

**Figure 2:**
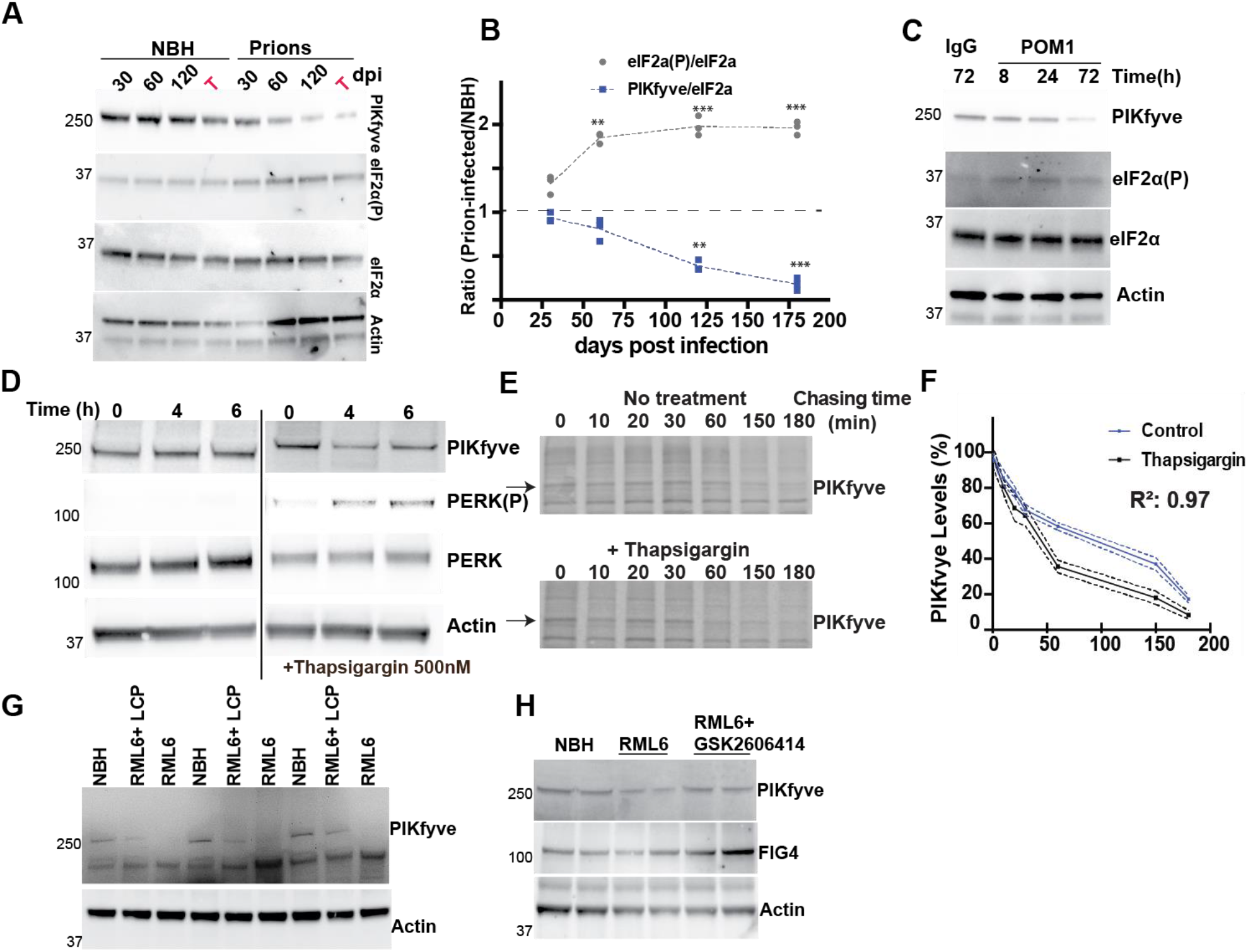
ER stress regulates PIKfyve expression levels. **A**: transient PIKfyve suppression in thapsigargin-treated Gt1 cells with recovery after 6 hours. PERK phosphorylation confirmed ER stress. **B**: Time course of eIF2α phosphorylation in prion-inoculated C57BL/6 mice. T: terminal disease. **C**: PIKfyve/eIF2α and eIF2α(P)/eIF2α ratios assessed from Western blots as in B (n=3/group/time point). PIKfyve levels gradually decreased after 60 dpi whereas eIF2α became phosphorylated from 30 dpi until terminal disease. eIF2α phosphorylation and pPIKfyve depletion in POM1-treated *tg*a*20* COCS. Quantification in Figure S2E. **E**: Gt1 cells were treated with thapsigargin (0.5 µM, 4 hours) or DMSO for control, and metabolically pulsed with [^35^S]Met/Cys for 20 min at 37 °C. Cells were either immediately lysed or chased without ^35^S at 37 °C for ≤3 hours. Proteins were immunoprecipitated with antibodies to PIKfyve and subjected to SDS-PAGE and autoradiography. **F**: Autoradiographic signals expressed as percentage of the initial PIKfyve levels. Thapsigargin shortened the half-life of PIKfyve from 79 to 34 min. First-order kinetics was assumed. Dotted lines: standard deviation. **G**: Prion-infected *tg*a*20* mice were treated with the polythiophene LIN5044 (LCP) or vehicle. LCP partially restored PIKfyve levels (quantification: Figure S2E). **H**: *tg*a*20* COCS were inoculated with RML or NBH, optionally treated with GSK2606414 starting at 21 dpi, and lysed at 35 dpi. Depletion of PIKfyve in prion-infected samples was largely rescued by GSK2606414 (quantification: Fig. S2G)

If PIKfyve depletion mediates prion neurotoxicity, it may be attenuated by antiprion therapeutics. The conjugated polythiophene LIN5044 binds PrP^Sc^, antagonizes prion replication^20,21^, delays prion pathogenesis in vivo and reduces spongiosis^22^. Remarkably, prion-infected mice treated with LIN5044 not only showed prolonged survival but also partially restored PIKfyve levels (Fig. 2G, S2E). We then administered GSK2606414, an inhibitor of the protein kinase RNA-like endoplasmic reticulum kinase (PERK) arm of the UPR^19^, to prion-infected *tg*a*20* COCS at 21 dpi (Fig. S2F). GSK2606414 partially restored PIKfyve levels (Fig. 2H, S2G), strengthening the notion that chronic ER stress is the direct cause of PIKfyve downregulation.

Dynamic acylation and deacylation control protein stability^23^ and can be affected by the UPR^24^. Being juxtaposed to endosomal membranes, PIKfyve is a plausible candidate for acylation. Acyl-resin assisted capture (acyl-rac) of mouse brain homogenates showed that PIKfvye, but not the cysteine-less translocon-associated protein-α (TRAPα), was acylated (Fig. 3A). Prion-infected samples, however, showed massively reduced PIKfyve acylation already at 60 dpi, a time point showing vigorous UPR but only minimal depletion of total PIKfyve (Fig. 2A, 3B, S3A).

**Figure 3:**
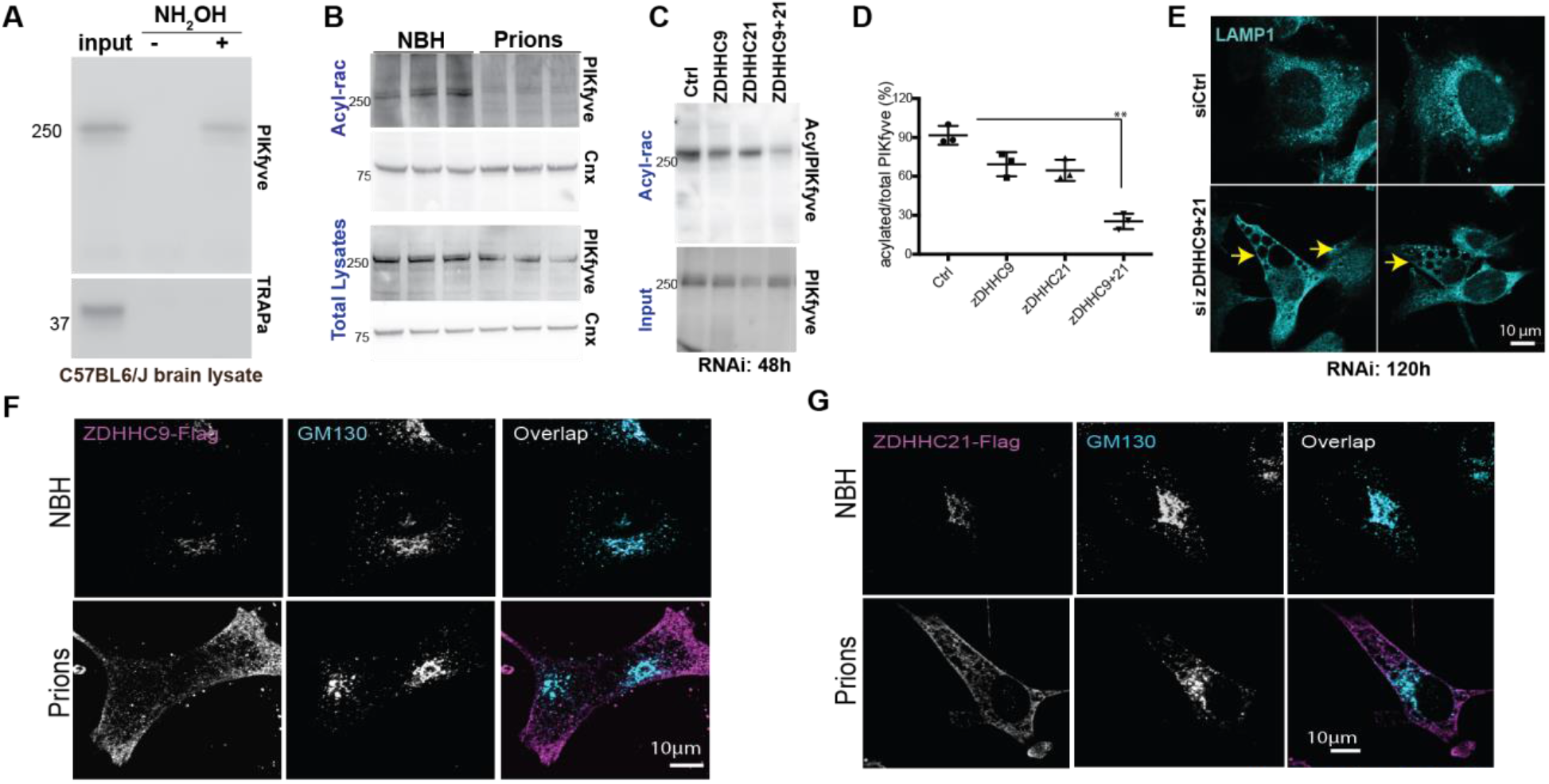
Prion infection causes deacylation and degradation of PIKfyve. **A:** Acyl-rac captured hydroxylamine-cleavable, acylate PIKfyve from mouse brains. Control: TRAPα, a non-acylated protein. Input: 10% of lysate. **B:** Brain lysates from prion-inoculated mice (60 dpi) precipitated by Acyl-rac. PIKfyve, but not calnexin (Cnx), was deacylated in prion-infected samples. Each lane represents an individual mouse. Lower panel: lysate used for Acyl-rac. **C-D:** Acyl-rac of Gt1 cells subjected to siRNA against zDHHC9, zDHHC21 or both (72 hours post transfection), showing synergistic decrease of PIKfyve acylation by suppression of zDHHC9 and 21 (quantified in D). Each dot represents a separate experiment (unpaired t-test). **F:** Gt1 cells were transfected with siRNA against zDHHC9+21 and stained with LAMP1. Arrows: vacuoles. **F-G**: Flag-tagged zDHHC9 or zDHHC21 were expressed in Gt1 cells were transfected with plasmids encoding. After 72 hours, zDHHC9/21 (magenta) were localized to the Golgi apparatus (GM130^+^, cyan) in NBH-exposed cells but not in prion-infected cells.

An acylation prediction database^25^ pointed to cysteine residues 202 and 203 as potential palmitoylation sites in all three isoforms of PIKfyve (Fig. S3B). We therefore expressed PIKfyve-GFP or PIKfyve^C202+203A^GFP, in which both residues were replaced by alanines, in Gt1 cells. Acyl-rac confirmed that mutagenesis of cysteine residues 202 and 203 prevented PIKfyve acylation (Fig. S3C). We next investigated the function of the acylation machinery in prion infections. The mouse genome encodes 23 acyl transferases characterized by a zinc-finger-domain and an aspartate-histidine-histidine-cysteine tetrapeptide (zDHHC), denoted zDHHC1-24. Quantitative real-time PCR (qPCR) on terminally scrapie-sick mouse brains failed to identify altered mRNA levels of any zDHHC member (Fig. S3D). We then mined a database of transcriptional changes during the course of prion infection^17^. None of the zDHHC mRNAs were altered at any stage of the disease (Fig. S3E). Next, we suppressed each zDHHC individually by siRNA. All zDHHC mRNAs were reduced by >80%. After 72 hours, cells were harvested and subjected to acyl-rac. Only the knockdown of zDHHC9 and zDHHC21 modestly reduced the acylation of PIKfyve (Fig. 3D, S3F). We next knocked down zDHHC9 and 21 individually vs. simultaneously in Gt1 cells. Acyl-rac at 48 hours showed synergy of the zDHHC9/21 siRNA mix (Fig. 3C-D). Crucially, at 120 hours the zDHHC9/21 siRNA mix induced not only a downregulation of PIKfyve but also the appearance of vacuoles (Fig. S3G, 3E) similar to those of prion-infected cells.

Next, we expressed in Gt1 cells zDHHC9 and zDHHC21 fused to a C-terminal flag-tag. While zDHHC9 and zDHHC21 localized to Golgi complexes of NBH-treated cells, in prion-infected cells both enzymes became drastically mislocalized to the cell periphery (Fig. 3F-G). The mislocalization of zDHHC9/21 may explain the loss of PIKfyve acylation and its degradation thereby driving spongiogenesis.

## Cellular consequences of prion-induced spongiosis

Since PIKfyve is necessary for the maturation of late endosomes into lysosomes, prion-associated spongiosis may result from a blockade of endosomal maturation. We found that LAMP1, a marker for late endosomes/lysosomes, was associated with vacuoles in both prion-infected Gt1 cells and in Gt1 cells depleted of PIKfyve using shRNA (Fig. 4A). We wondered whether the association of LAMP1 with vacuoles can also be observed in other model systems of prion disease. We infected COCS generated from 9-day old *tg*a*20* mice with RML. At 50 dpi, we detected LAMP1 upregulation in neurons (Fig. 4B, S4A). As in Gt1 cells, vacuoles in COCS were lined by LAMP1 (Fig. 4C). Treatment of *tg*a*20* COCS with POM1 (10 days) also resulted in microvacuolation and LAMP1 in neurons (Fig.4D, S4B). These results suggest that spongiosis originates from late endosomal/lysosomal compartments. To differentiate between these two possibilities, we stained PIKfyve-depleted Gt1 cells (72 hours after siRNA transfection) with the ratiometric pH probe Lysosensor Yellow/Blue (5 µM, 5 min). Flow cytometry revealed no enhanced acidification in vacuolated cells (Fig. 4E), suggesting that vacuoles represent stalled prelysosomal compartments.

**Figure 4:**
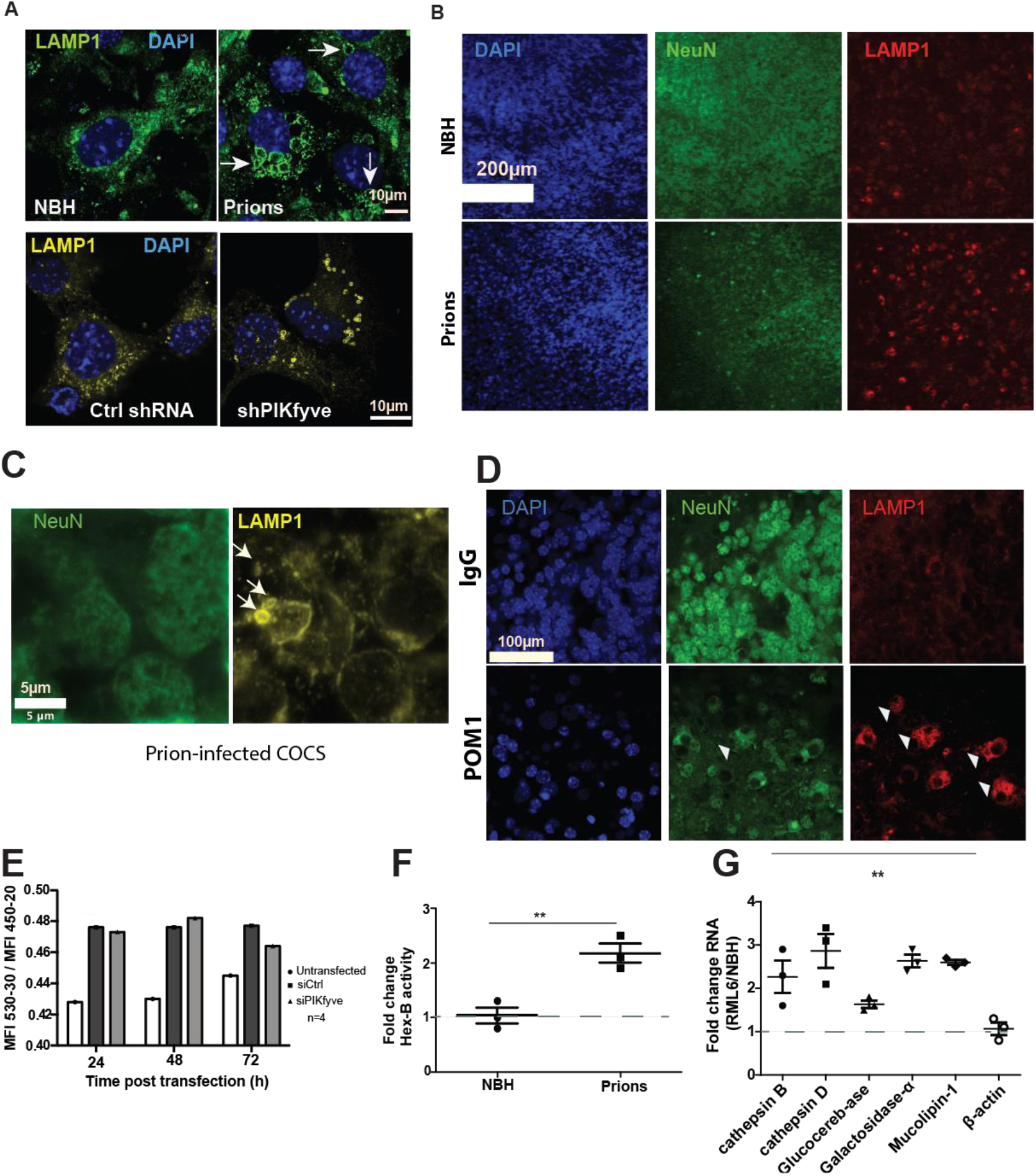
Loss of PIKfyve induces lysosomal defects. **A:** Upper row: prion-infected Gt1 cells (75 dpi) immunostained for LAMP1 showing prominent vacuoles. Control: NBH-inoculated cells. Lower row: Gt1 cells transfected with shRNA targeting PIKfyve or luciferase (control) for 5 days and immunostained for LAMP1. Depletion of PIKfyve resulted in LAMP1^+^ vacuoles (yellow). Nuclei were stained by DAPI. **B:** Prion-infected *tg*a*20* COCS (45 dpi) immunostained with NeuN and LAMP1. Control: NBH-exposed COCS. Nuclei: DAPI. Prion infection induced LAMP1 upregulation in neurons (quantification: Fig. S4A). **C:** Higher magnification showing LAMP1^+^ vacuoles (arrowheads) in prion-infected COCS. **D:** *tg*a*20* COCS treated with POM1 or control IgG (10 days) and immunostained with NeuN and LAMP1. Nuclei: DAPI. POM1-treated COCS showed increased LAMP1 expression (quantification: Fig. S4B). NeuN^+^ cells exhibited vacuoles (arrowheads). **E:** Gt1 cells were transfected with control siRNA (scrambled) or siRNA targeting PIKfyve for up to 72 hours and stained with LysoSensor Yellow/Blue. Cells were gated based on emission spectra, and mean fluorescence intensities (MFIs) per sample were quantified. Increased 530/450 MFI ratio indicates enrichment of acidic compartments. No acidic compartment expansion in PIKfyve-depleted cells. **F:** β-hexosaminidase A activity in brain lysates from terminally scrapie-sick and NBH-inoculated mice. Control: NBH inoculated mice. Prion-infected brain lysates showed 2-2.5 fold increased activity. Panels show independent triplicates. Statistics: unpaired t test. **G:** Lysosomal enzymes, but not β-actin, were elevated in brains of terminally scrapie-sick mice. Panels show independent triplicates. Statistics: ANOVA.

Increased activity of LAMP1 and other lysosomal enzymes is a frequent feature of lysosomal dysfunction. Indeed, we found a three-fold increase in the enzymatic activity of lysosomal hexosaminidase-β in brains of prion-infected mice (Fig. 4F). In addition, we monitored the RNA expression levels of cathepsin-D, cathepsin-A, glucocerebrosidase, α-galactosidase and mucolipin-1, previously shown to be upregulated in lysosomal diseases^26^, and found all of them to be upregulated in brains of terminally scrapie-sick mice (Fig. 4G). Lysosomes undergo cycles of fission and fusion, which are impaired by the absence of PIKfyve, resulting in coalesced endolysosomes^27^. The consequences include nuclear translocation of TFEB and depletion of TRPML1, a channel controlling lysosomal size^28-30^. TRPML1, whose lysosomal association relies on PIKfyve^31^ and was abolished by shRNA-mediated PIKfyve knockdown in Gt1 cells (Fig. S4C), was co-precipitated by anti-LAMP1 antibodies in NBH-inoculated but not in scrapie-sick mouse brains (Fig. S4D). Total TRPML1 protein levels were unaltered.

Upon dephosphorylation, transcription factor EB (TFEB) translocates to the nucleus and promotes coordinated expression of lysosomal genes^26,32,33^. TFEB became progressively dephosphorylated in prion-infected mouse brains whereas total TFEB levels remained constant (Fig. 5A). We then examined a subset of TFEB-regulated lysosomal genes in terminally prion-sick brains. The mRNA levels of all investigated genes were conspicuously increased, whereas the non-lysosomal gene STAT3 was not (Fig. 5B). These findings could reflect the vivacious proliferation of activated microglia in terminal prion disease. We therefore investigated the lysosomal status of Gt1 cells (Fig. 5C) and COCS (Fig. 5D). We found that prion infection led to TFEB dephosphorylation in both models. Moreover, prion-infected Gt1 cells (75 dpi) displayed nuclear translocation of TFEB (Fig. 5E) and prion-infected Gt1 cells (75dpi), as well as Gt1 cells depleted of PIKfyve by shRNA, showed upregulation of TFEB responsive genes (Fig. 5F-G). This disproves that lysosomal gene upregulation merely indicates distorted tissue composition, and suggests that TFEB-mediated reactions are a cell-autonomous consequence of PIKfyve depletion. Finally, we tested the relationship between TFEB and lysosomal homeostasis by treating prion-infected Gt1 cells with siRNA against TFEB. 72 hours post siRNA transfection, the expression of TFEB-responsive genes was normalized, yet the vacuolation of prion infected cells was unaltered (Fig. S4E-G), suggesting that TFEB-related lysosome pathologies may be a consequence rather than a cause of spongiosis.

**Figure 5:**
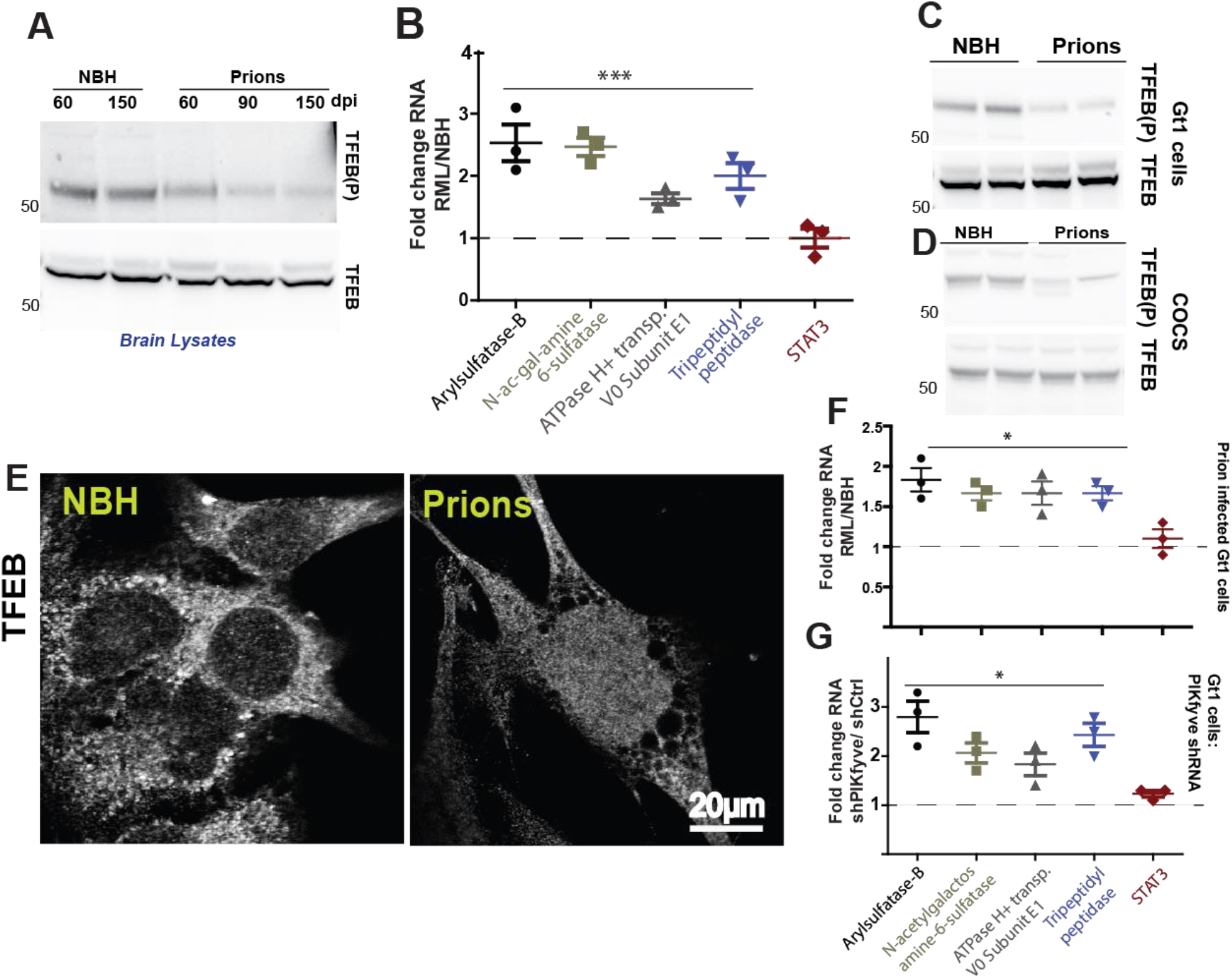
Prions induce lysosomal dysregulation via TFEB. **A:** Time course of TFEB phosphorylation in prion-infected C57BL/6 mice. Total TFEB was unchanged but reduced TFEB phosphorylation was evident at ≥90 days. **B:** TFEB-controlled transcripts in brains of terminally scrapie-sick C57BL/6 mice were measured by qPCR. Prion infection led to upregulation of these genes, but not of STAT3. Control: NBH-inoculated mice. ANOVA on independent triplicates **C:** Prion-infected Gt1 cells (75dpi) showed reduced TFEB phosphorylation. Total levels of TFEB remain unchanged. **D:** COCS prepared from *tg*a*20* mice were infected with RML prions and cultured for 5 weeks. Cell lysates were prepared at 4 wpi followed by Western blot analysis using anti-phospho-TFEB antibody. Activated (dephosphorylated) TFEB was increased in RML infected COCS. Total levels of TFEB remain unchanged. **E:** Gt1 cells (as in **D**) were fixed and stained with anti-TFEB antibody. Prion-infected cells showed nuclear translocation of TFEB. **F:** TFEB responsive genes were upregulated in Gt1 cells chronically infected with RML prions (75 dpi). Panels depict independent triplicates. Statistics: ANOVA. **G:** TFEB responsive genes were upregulated in cells transfected with shPIKfyve for 5 days compared to shCtrl (shRNA against luciferase). Panels depict independent triplicates. Statistics: ANOVA.

The following hierarchy of events emerges from these findings. Chronic activation of the UPR leads to mislocalization of acyl transferases. This, in turn, compromises PIKfyve acylation, reduces its half-life, and destabilizes its association with VAC14 and FIG4. The resulting reduction of PI(3,5)P_2_ stalls lysosome maturation and induces inappropriate TFEB-dependent transcriptional responses. We challenged this model by interfering with each of its predicted checkpoints. Firstly, we asked whether overexpression of zDHHC9 and zDHHC21 might restore PIKfyve levels and abrogate vacuole generation in prion-infected Gt1 cells (75 dpi). Indeed, transfection (72 h) with both zDHHC9 and zDHHC21 partially restored PIKfyve levels and reduced the number of vacuolated cells (Fig. 6A-B). Secondly, we attenuated the UPR in prion-infected Gt1 cells (75 dpi) by lentiviral transduction of GADD34, which dephosphorylates eIF2α^34^. Again, we found restoration of PIKfyve levels and a significant decrease in the frequency of vacuolated cells (Fig. 6C-D). Besides confirming that increasing PIKfyve levels suffices to prevent vacuolation, these results establish the directionality of chronic UPR, delocalization of acyl transferases and PIKfyve depletion. Thirdly, we reasoned that prion-induced spongiosis could be repressed by the PIKfyve adduct, PI(3,5)P_2_. We treated prion-infected *tg*a*20* COCS for 45 days with bodipy-PI(3,5)P_2_ (bPIP), a fluorescent water-soluble analog of PI(3,5)P_2_ (5µg/ml). Infection with RML led to complete ablation of cerebellar granule layer (CGN), which was attenuated by bPIP (Fig. 6E). Furthermore, bPIP reduced the occurrence of LAMP1^+^ vacuoles in prion-infected COCS, and restored the homeostasis of TFEB-responsive genes (Fig. S5A-B). Next, we treated *tg*a*20* COCS with POM1 for 14 days in the presence or absence of bPIP. The subsequent ablation of the CGN was again attenuated by bPIP (Fig. 6F). We then treated prion-infected Gt1 cells with bPIP (70 dpi, 20 µg/ml). After 12 hours, >60% of cells incorporated bPIP into lysosomes (Fig. S5C-D). Fresh bPIP, or water for control, was added every 12 hours over 72 hours (Fig. S5E). bPIP reduced the number of vacuolated cells (Fig. 6G) and restored the homeostatic expression of TFEB responsive genes (Fig. S6A), but did not alter the accumulation of PrP^Sc^ (Fig. S6B). Hence bPIP acts downstream of PrP^Sc^ generation. To further probe the specificity of bPIP, we depleted PIKfyve from Gt1 cells using shRNA. The ensuing vacuolation was conspicuously reduced by bPIP (Fig. S6C).

**Figure 6:**
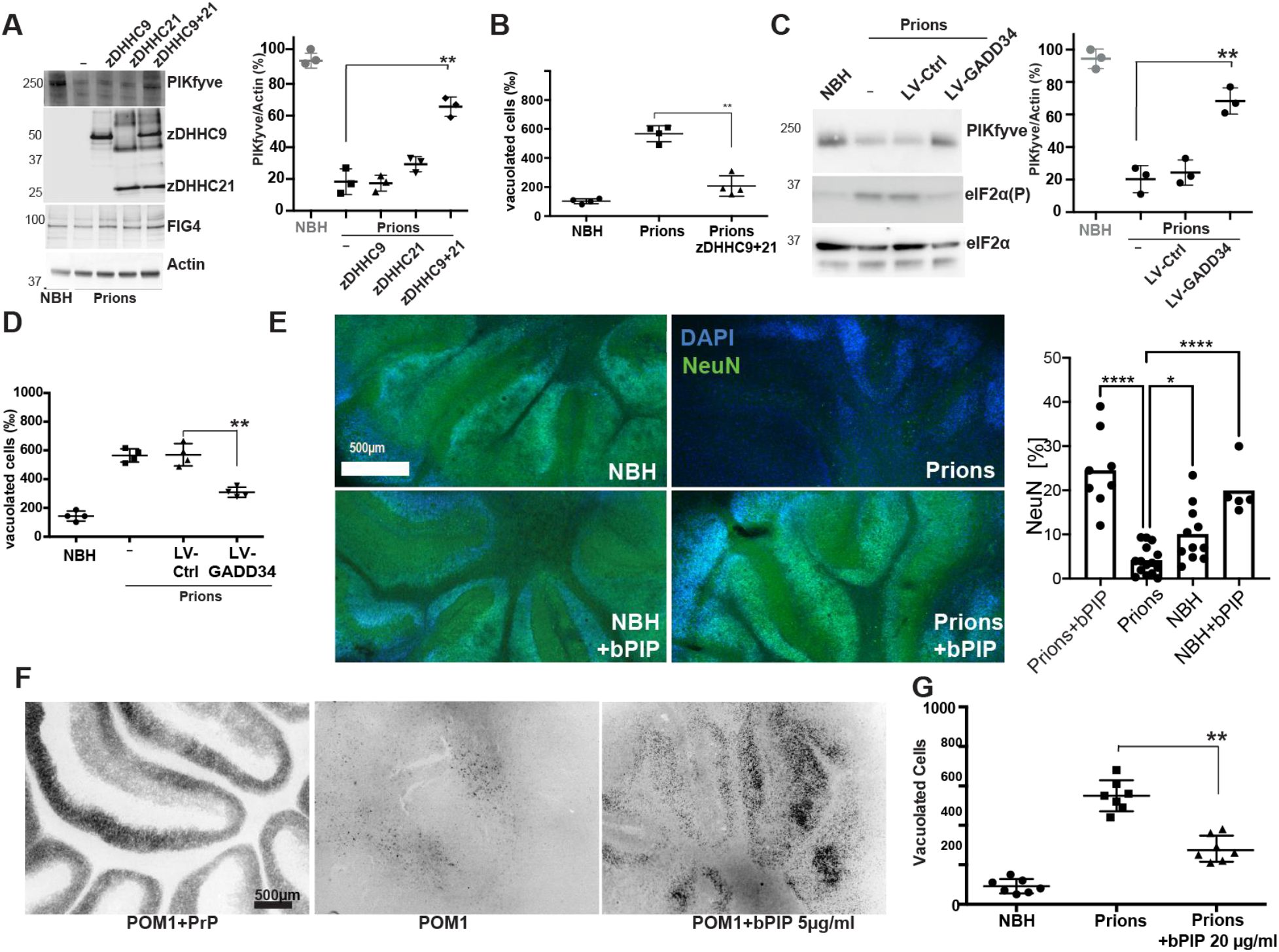
PI_(3,5)_P2 attenuates spongiosis and lysosomal dysfunction. **A:** Transient overexpression of Flag-tagged zDHHC9+21 (72 hours) restored PIKfyve levels in prion-infected Gt1 cells. Right: Western blot quantification. Here and henceforth, each dot represents an individual experiment (unpaired t-test). **B:** Cotransfection of zDHHC9+21 reduced vacuolation in prion-infected Gt1 cells. Each dot represents a separate experiment (1000 cells/experiment, χ^2^: *p*<0.001). **C:** Prion-infected Gt1 cells were lentivirally transduced with GADD34 at 75 dpi. Control: empty lentiviral vector. At 79 dpi, GADD34 overexpression had normalized eIF2α phosphorylation and restored PIKfyve levels. Right: Quantification of western blots (unpaired t-test). **D:** Number of vacuolated cells from the experiment shown in C. GADD34 expression rescued the total number of vacuolated cells (1000 cells/experiment, χ^2^: *p*<0.001). **E:** *tg*a*20* COCS were infected with RML and optionally treated with bPIP (5µg/ml). At 45 dpi, NeuN morphometry revealed ablation of cerebellar granule layer (CGN) in prion infected slices and its rescue by bPIP. Control: NBH treated COCS. Each dot represents an individual slice (Statistics: ANOVA) **F:** *tg*a*20* COCS were treated with POM1 (optionally pre-blocked with recPrP) and treated with bPIP (5µg/ml). At 14 dpi NeuN morphometry revealed POM1-induced ablation of cerebellar granule layer (CGL) and rescue by bPIP. **G:** Prion-infected Gt1 cells were treated with bPIP for 3 days. The number of vacuolated cells was reduced (1000 cells/experiment, χ^2^: *p*<0.001). h

The above evidence shows that neuronal vacuolation is tightly coupled to PIKfyve depletion in disparate experimental models, and the virulence of human prion diseases correlates with the extent of PIKfyve depletion. Prion infection causes a sustained UPR, as do other neurodegenerative diseases^35^ including granulo-vacuolar degeneration of Alzheimer’s disease^36^ and inclusion-body myopathy which features prominent myofiber vacuolation^37^. This begs the question whether PIKfyve may play a role in vacuolating diseases other than prion infections.

## Acknowledgements

We thank Rita Moos and Petra Schwarz for technical help, Dr. Sebastian Jessberger for donating the zDHHC9 and zDHHC21 plasmids, Dr. Giovanna Mallucci for providing the lentivirus overexpressing GADD34, Dr. Pamela Mellon for Gt1 cells, Dr. Claudia Scheckel for critical comments on the manuscript and assistance in writing and members of Aguzzi lab for their critical inputs on the manuscript. The cartoon depicting the human brain was obtained from The Database Center for Life Science and licensed under Creative Commons Attribution-Share).

## Author contributions

AA and AKKL designed the experiments and wrote the manuscript. AA initiated and supervised the project, and wrote the manuscript. AKKL performed, or contributed to, all experiments including western blots, immunoprecipitations, acylation assays, q-PCR assays, experiments on cerebellar organotypic cultured slices (COCS), immunofluorescence, imaging, vacuolation rescue experiments. EL performed imaging on prion-infected cells, ER stress experiment in GT1 cells and alternative splicing of PIKfyve. KF contributed to the western blots on human CJD samples, performed the bodipy-PI(3,5)P_2_ rescue experiments on RML treated and POM1-treated slices, performed quantifications on slice cultures and wrote the bioethical application to perform the experiments on human samples. UH contributed to the preparation of COCS, prepared the COCS for the rescue of PIKfyve levels upon treatment with GSK2606414, treated the mice with Lin5044 LCP and generated the brain lysates, and generated shRNA against PIKfyve. ML performed the FACS experiment to monitor the content of the vacuoles. RM set up conditions for performing western blots using various antibodies generated the prion-infected Gt1 cells, performed western blots, IP, staining of infected Gt1 cells, contributed to the bodipy-PI(3,5)P_2_ rescue experiment in Gt1 cells and performed immunofluorescence on prion-infected cells and their decontamination for imaging. All authors approved the final version of the manuscript.

## Data availability

All original data has been included in the article. Raw data and reagents will be made available by the corresponding author upon request.

