## Supplementary Figures and Methods for "Loss of PIKfyve drives the spongiform degeneration in prion diseases"

**Supplementary Information**

**Figure S1**

**A:** Liver and spleen lysates were prepared from terminally sick C57BL/6 mice infected with RML6 prions and, for control, age-matched mice injected with NBH. Samples (50 µg) were subjected to SDS-PAGE followed by Western blot with antibodies against PIKfyve and actin. In both organs PIKfyve levels were unaffected by prion infection. **B:** Brain lysates were prepared from terminally scrapie-sick *tga20* mice (overexpressing $PrP^C$) infected with RML6 prions. For control we used age-matched mice inoculated with NBH. Western blot analysis revealed downregulation of PIKfyve but no effect on FIG4. Actin: loading control. **C:** Brain lysates were prepared from terminally sick C57BL/6 mice infected with RML6 prion. For control we used age-matched mice inoculated with NBH. Samples (50 µg) were subjected to SDS-PAGE followed by Western blot with antibodies against PIKfyve, FIG4, VAC14, Tsg101, SOD2, MGRN1 and actin. Only PIKfyve was significantly downregulated in prion infection ($p<0.001$); Statistics: ANOVA. **D:** Gt1 cells infected with RML6 prions or NBH were treated with siRNA against $PrP^C$ for 96h at 70dpi. Phase contrast microscopy revealed a significant decrease in the number of vacuolated cells upon depletion of $PrP^C$ in RML6 infected cells. 1000 cells were manually quantified for the presence of vacuoles. Statistics: Chi-Square test. **E:** Electron microscopy image showing membrane-lined microvacuoles (arrows) in prion-infected COCS at 56 dpi. The vacuoles contained degenerating organelles and other cellular debris. Yellow arrows: vacuoles. **F:** Cerebellar organotypic cultured slices (COCS) were generated from *tga20* mice pups and inoculated with RML6 or, as a control, with NBH. At 45 dpi, when neurodegeneration was prominent, COCS were lysed and equal amounts of the protein was used for western blot analysis. PIKfvye was depleted in RML6 infected COCS. Loading control: actin. Right: Quantification of the western blot shows a significant reduction in PIKfyve levels. Experiments were repeated three times independently. **G-H:** Representative hematoxylin and eosin (H&E) staining of the human cortical brain tissue obtained at autopsy from patients who suffered from Type-1 or Type 2-CJD. AMmore extensive vacuolation can be observed in Type-2 CJD. **I:** Gt1 cells were transfected with siRNA against VAC14 or FIG4. For control we used an siRNA containing a scrambled sequence. Cell were harvested 72 hours post transfection and 50 µg of protein was migrated on SDS-PAGE followed by Western blotting using anti PIKfyve, FIG4 and VAC14 antibodies. PIKfyve levels were reduced in cells with VAC14 and FIG4 downregulation. **J:** The $\log_2$ fold changes in the mRNA expression of PIKfyve, FIG4 and VAC14 during the course of prion disease were extracted from a longitudinal database of transcriptional changes in prion disease progression [1]. None of the genes show significant alterations during the progression of the prion disease. **K:** cDNA generated from the lysates described in Fig. S1E was subjected to qPCR using primers targeting PIKfyve. No change was observed in the expression levels of PIKfyve after prion infection. **L:** Total RNA was isolated from COCS treated for 72 hours with POM1 or IgG for control. As a further control we used POM1 preincubated with recombinant PrP (thereby blocking its paratope). RNA was retrotranscribed, and cDNA was subjected to qPCR using primers against PIKfyve. PIKfyve RNA levels remained unaltered after treatment with POM1. **M:** Genomic map of the three differentially spliced isoforms of PIKfyve. Vertical lines denote the exons (not to scale). Variant 1 represents the protein coding and the major isoform and consists 42 exons. Variant 2 and 3 are the splice variants which contain exons which result from differential splicing (marked in red). Specific primers (marked in blue) were used spanning the exons that are specific to each of the isoform measure their levels in brains using real time quantitative PCR (qPCR). **N:** Total RNA was isolated from the brains of terminally sick C57BL/6 mice infected with RML6 prions and as a control RNA was isolated from NBH infected mice (same samples as in D). 1 µg of RNA was subjected to reverse transcription to generate cDNA followed by real time quantitative PCR (-PCR) using primers targeting each of the three isoforms of murine PIKfyve individually. Variant 1 represents the full length and is predominant isoform expressed in the brains. Variant 2 and 3 are expressed in low amounts in comparison to variant 1. The levels of all the isoforms were unaffected by prion infection. Each dot represents an individual mouse; Statistics: ANOVA.

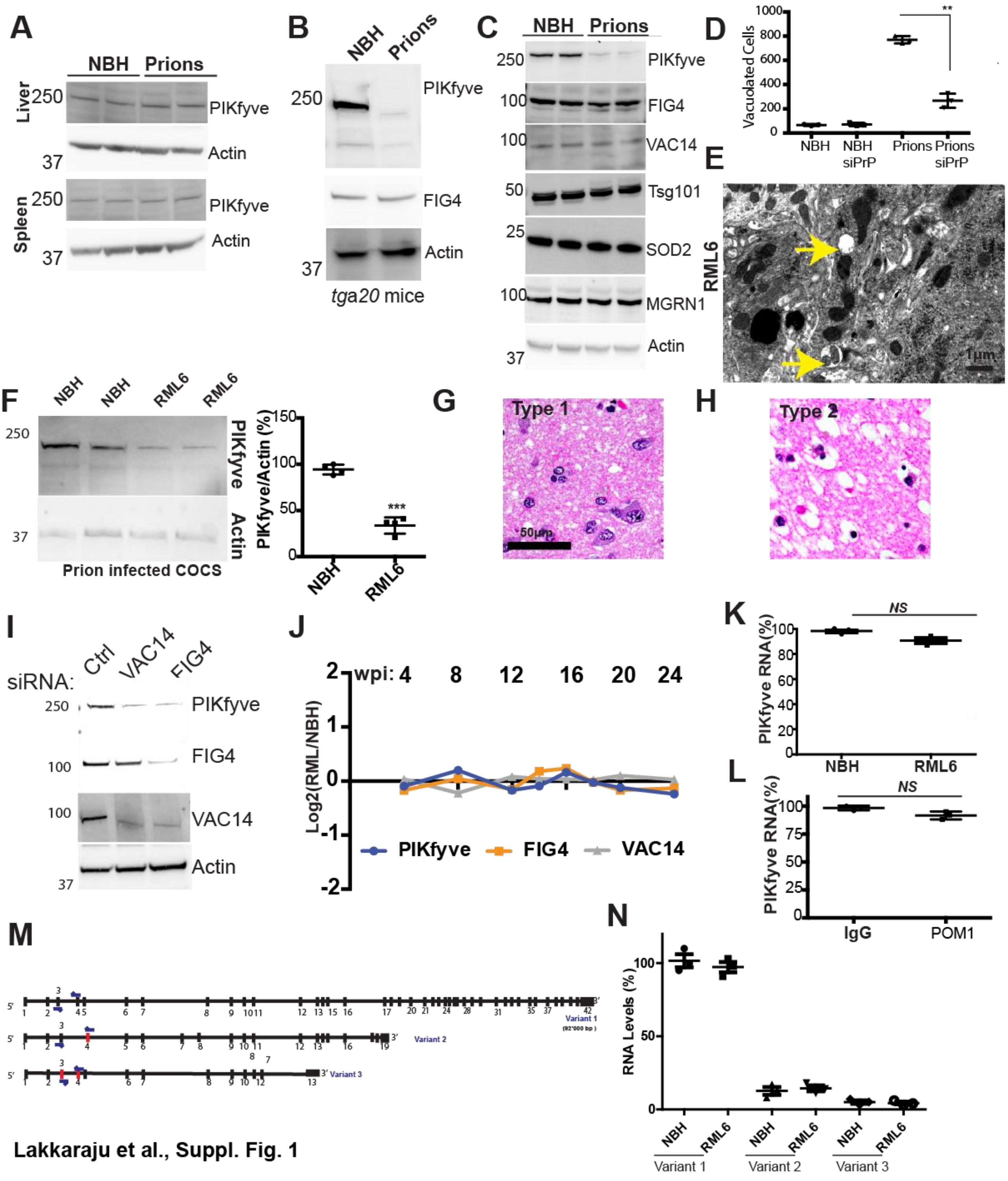


**Lakkaraju et al., Suppl. Fig. 1**

**Figure S2:**

**A:** Quantification of the western blot from Fig. 2D. Treatment of COCS with POM1, reduces the levels of PIKfyve starting 48h post treatment. Each dot represents an individual experiment. Statistics: Unpaired t-test. **B:** RNA was isolated from Gt1 cells treated with thapsigargin (0.5 µM for 4h), retrotranscribed and subjected to quantitative real-time PCR. PIKfyve mRNA remained unaltered. Panels depict three independent experiments. Statistics: Unpaired t-test. **C:** Quantification of western blot from Fig. 2D. The experiment was repeated three independent times. PIKfyve levels significantly decrease 4 hours post thapsigargin treatment but recover back at 6 hours post treatment. **D:** Gt1 cells were treated with thapsigargin (0.5µM for 4hours) followed by imaging using a phase contrast microscope at 20x magnification. **E:** Quantification of the western blot from Fig. 2G. Each lane in the western blot contained lysates from an individual mouse. PIKfyve levels are partially restored in the presence of Lin5044. Statistics: Unpaired t-test. F**G:** Schematics of GSK2606414 treatment of prion infected COCS. Prion-infected COCS were continuously treated with GSK2606414 starting from day 21 until day 45 when the samples were lysed, and protein and RNA was harvested for further analysis. **G:** Quantification of western blot from Fig. 2H. Treatment of COCS with GSK2606414 attenuated the loss of PIKfyve. Each dot represents an individual experiment. Statistics: Unpaired t-test.

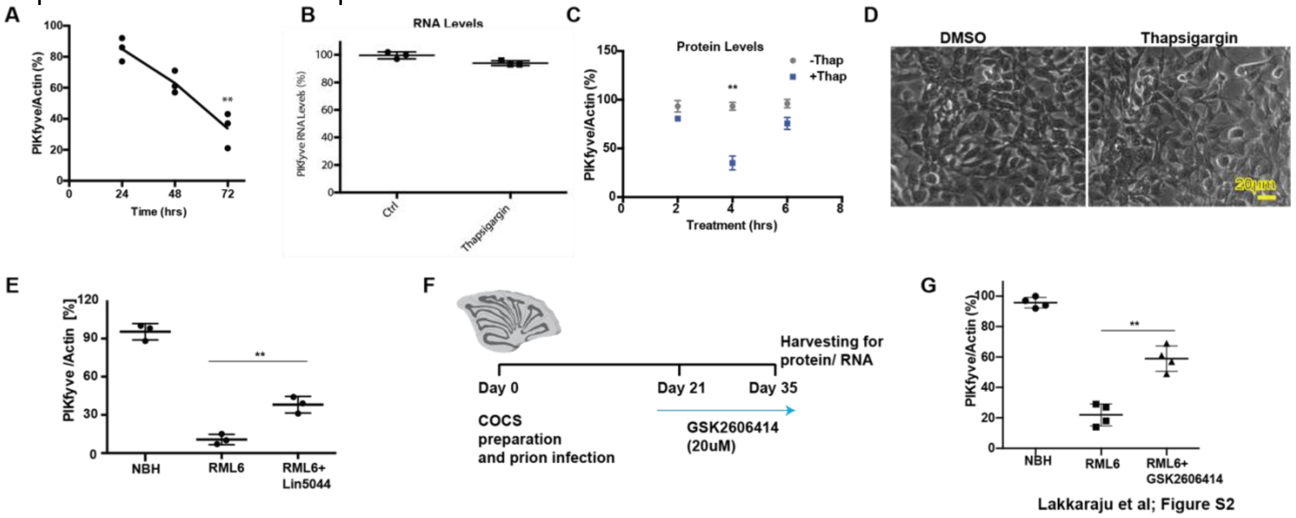

**Figure S3**

**A:** Quantification of western blot from Fig. 3B. Acylated PIKfyve levels were significantly downregulated in prion infected mice at 60 dpi. Each dot represents an individual mouse. Statistics: Unpaired-t test. **B:** Sequence alignment of all the three isoforms of PIKfyve revealed the presence of conserved cysteines at positions 203 and 203, which were predicted by the acylation predictor to be the potential palmitoylation sites in PIKfyve. **C:** Gt1 cells were transiently transfected with full length wild type PIKfyve cDNA (PIKfyve$^{WT}$) or with PIKfyve plasmid where the cysteines at position 202/203 were replaced with alanines (PIKfyve$^{C202/203A}$ ). 48h post transfection, cell lysates were subjected to acyl-rac and immunoblotted with anti-GFP antibody. PIKfyve$^{C202/203A}$ was not acylated (right panel) and the total lysates revealed a lower expression of PIKfyve$^{C202/203A}$ (left panel). Calnexin was used an acylation control and as loading control. **DF:** Total RNA was isolated from the brains of terminally sick C57BL/6 mice infected with RML6 and subjected to reverse transcription. As control RNA from NBH infected mice was used. cDNA was subjected to qPCR using specific primers targeting each DHHC enzyme. No change in the levels of mRNA encoding any of the DHHC enzymes was observed between RML6-infected and NBH-exposed samples. **E:** The $\log_2$ fold changes in the mRNA expression of all zDHHC enzymes during the course of prion disease were plotted from the data available on transcriptional changes in prion disease progression[2]. None of the genes show significant alterations during the progression of the prion disease. **F:** Gt1 cells were transfected with siRNA against several zDHHC enzymes. After 72 hours, lysates were subjected to Acyl-rac and immunoblotted for PIKfyve. siRNA against zDHHC9 and 21 resulted in modest decrease in PIKfyve acylation. Lower panel represents the input (1/10$^{th}$ of the sample used from Acyl-rac). The experiment was repeated two times independently to confirm the identity of zDHHC responsible for acylating PIKfyve. **G:** Gt1 cells were treated with either a siRNA cocktail targeting zDHHC9/21 or a control siRNA (non targeting siRNA). 120 hours post transfection, cells were lysed and immunoblotted for PIKfyve. PIKfyve levels decreased in the prolonged absence of zDHHC9/21.

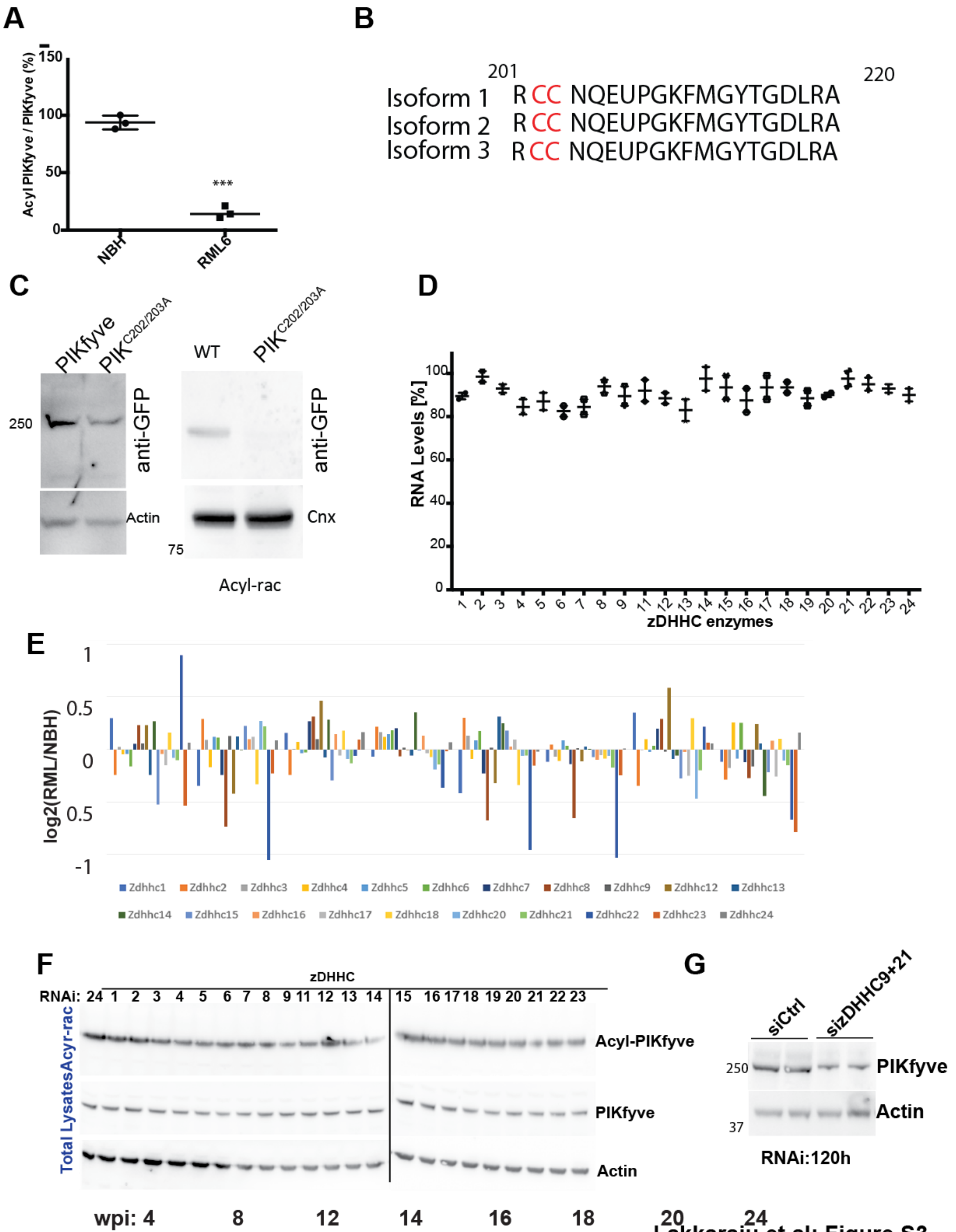


Lakkaraju et al; Figure S3

**Figure S4.**

**A-B:** Quantification of LAMP1 fluorescence in RML6 infected and POM1 treated *tga20* COCS. Only NeuN positive cells were considered for quantification. Statistics: unpaired t test. **C:** Gt1 cells treated with shRNA against PIKfyve for 5 days were fixed and stained with antibody against TRPLML1. Staining revealed a diffuse pattern of TRPML1 in PIKfyve-depleted cells whereas control cells displayed a typical lysosomal staining pattern. **D:** LAMP1 immunoprecipitates from prion-infected mouse brains. TRPML1 co-precipitated with LAMP1 in NBH-inoculated but not in prion-sick brains (left panel). Asterisk denotes heavy chain of the antibody. Right panel: total cell extracts indicate no change in the levels of TRPML1 in both NBH and prion-infected mouse brains. **E:** Gt1 cells with infected with RML6 and 70dpi, cells were treated with siRNA against TFEB for 5 days and were analyzed for lysosomal gene homeostatic levels using qPCR. TFEB downregulation resulted in restoration of lysosomal genes to their homeostatic levels. As control NBH treated cells were used. ARSB: Arylsulfatase B, GALNS: Galactosamine-6-sulfatase, ATP6V0E1: ATPase $H^+$-transporting-V0-Subunit-E1, TPP1: Tripeptidyl peptidase. Panels depict independent triplicates (Statistics: ANOVA). **F**: Phase contrast microscopy of cells in E. Vacuolation of prion-infected cells was not rescued by TFEB downregulation. Vacuoles: yellow arrows. **G:** Quantification of cells exhibiting vacuoles as a consequence of prion infection confirmed no rescue by TFEB downregulation. Each dot represents a separate experiment in which 1000 cells were counted. Statistics: Chi square test.

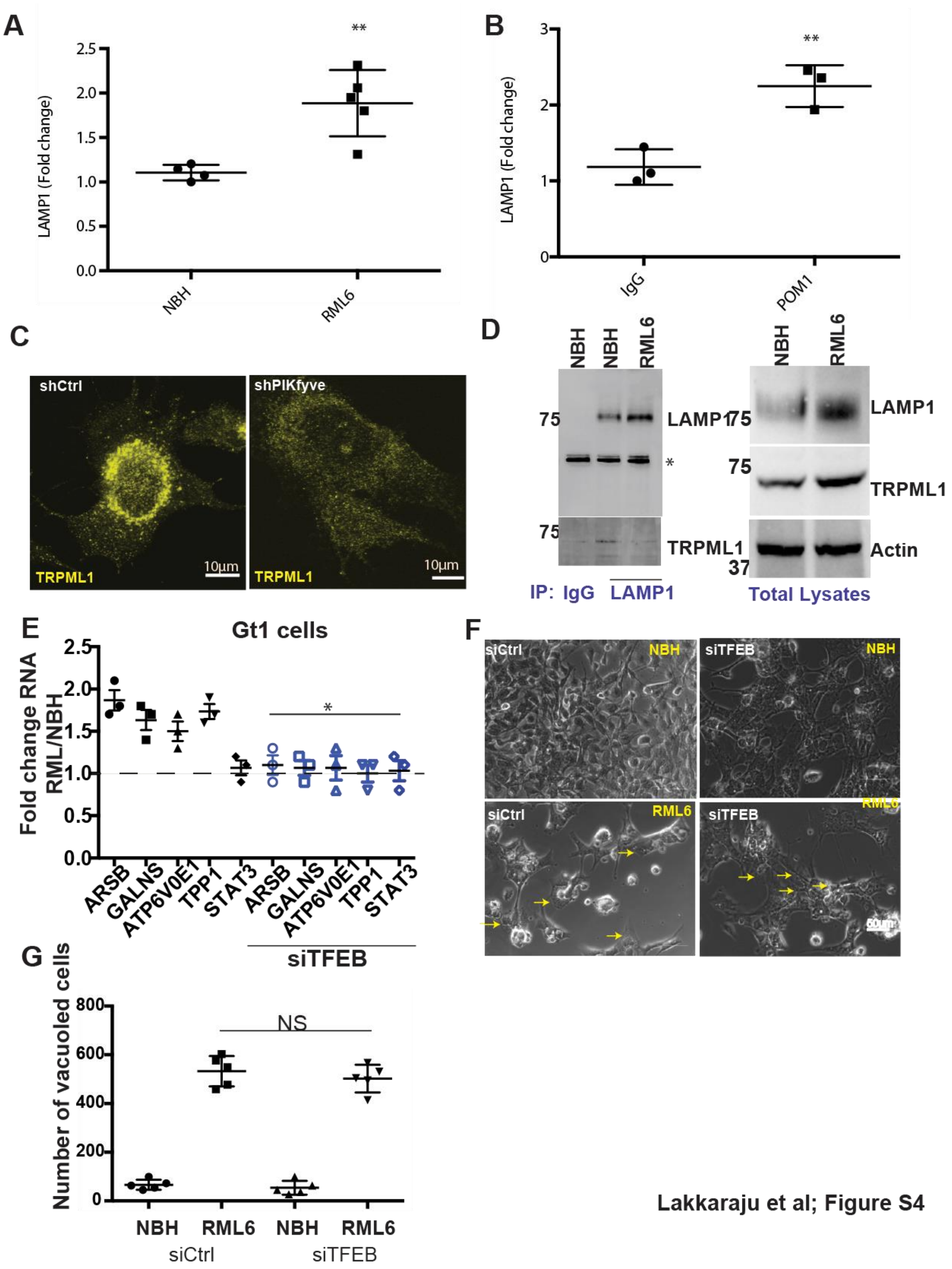


Lakkaraju et al; Figure S4

**Figure S5**

**A:** *tga20* COCS were infected with RML6 and optionally treated with bPIP (5µg/ml, 45 dpi) followed by immunostaining with LAMP1. NBH-exposed bPIP-treated COCS were used for control. Prion-infected COCS showed increased LAMP1$^{+}$ pixels; the increase was depressed by bPIP. Each dot represents an individual COCS. Statistics: ANOVA. **B:** COCS from A were analyzed for lysosomal gene expression by qPCR. bPIP treatment prevented the upregulation of TFEB-responsive genes in prion-infected COCS. ARSB: Arylsulfatase B, GALNS: Galactosamine-6-sulfatase, ATP6V0E1: ATPase H$^{+}$-transporting-V0-Subunit-E1, TPP1: Tripeptidyl peptidase. Panels depict independent triplicates (ANOVA). **C:** Gt1 cells were incubated with bPIP (10 µM; 12 hours) and subjected to cytofluorimetric analysis. 62% of cells took up bPIP. **D:** Gt1 cells were treated with bPIP as in A and incubated with Lysotracker (1 hour). bPIP localized to lysosomal regions stained by Lysotracker. **E:** Gt1 cells were infected with RML6 and treated with bPIP (10µM) for 5 days at 70 dpi. Depletion of PIKfyve resulted in vacuolation (left panel) which was significantly rescued by bPIP. Quantification shown in Fig. 6D; statistics: Chi-Square.

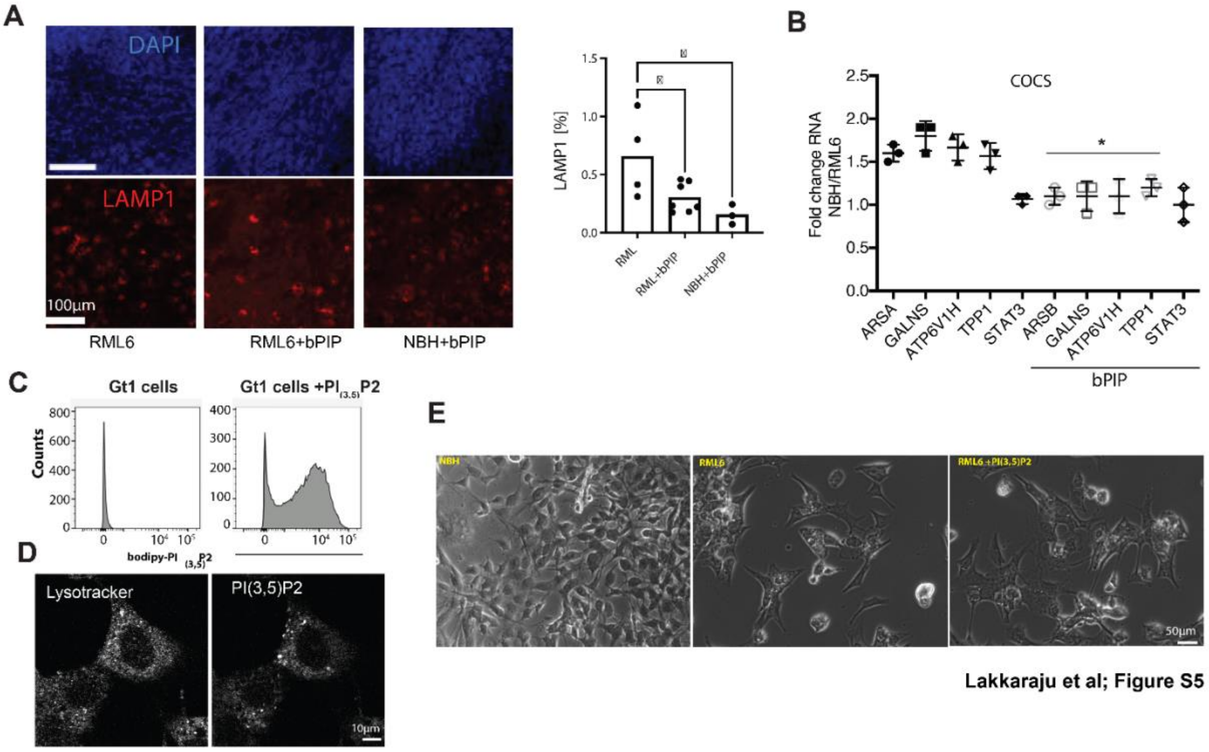

**Figure S6**
**A:** The Gt1 cell cultures depicted in Fig. 6G were analyzed for lysosomal gene expression by qPCR. bPIP treatment prevented the upregulation of TFEB responsive genes in RML6 prion infected Gt1 cells. ARSB: Arylsulfatase B, GALNS: Galactosamine-6-sulfatase, ATP6V0E1: ATPase $H^{+}$-transporting-V0-Subunit-E1, TPP1: Tripeptidyl peptidase. Panels depict independent triplicates (ANOVA). **B**: Representative Western blot from the prion infected Gt1 cells (Fig.6G) for total $PrP^{C}$ and proteinase K resistant $PrP^{Sc}$. The presence of $PrP^{Sc}$ in bPIP-treated cells suggests that bPIP acts downstream of prion aggregation. NBH-treated cells were used as controls. **C:** Gt1 cells were treated with shRNA against PIKfyve for 4 days in the presence or absence of bPIP (10 µM). 1000 cells were counted per experimental condition manually using phase contrast microscopy; each dot represents an individual experiment. Statistics: Chi-Square test.

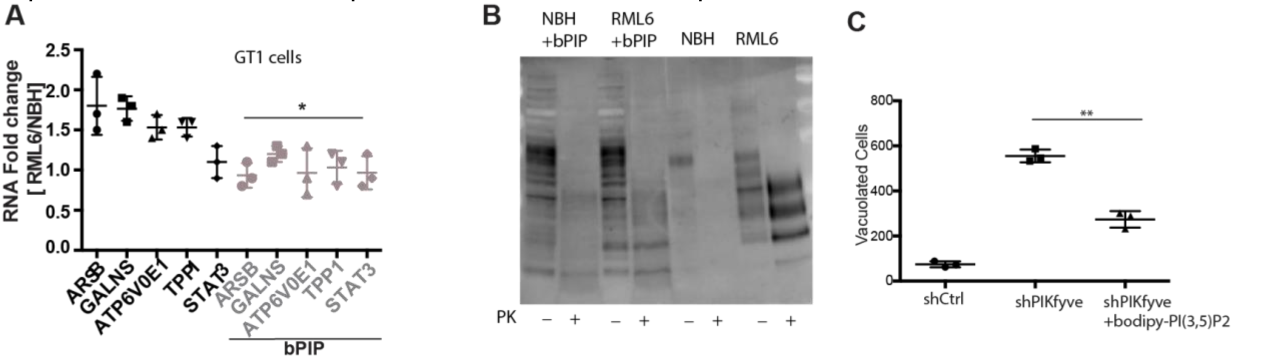


Lakkaraju et al; Figure S6

### Figure S7: Uncropped blots from the Manuscript

Figures

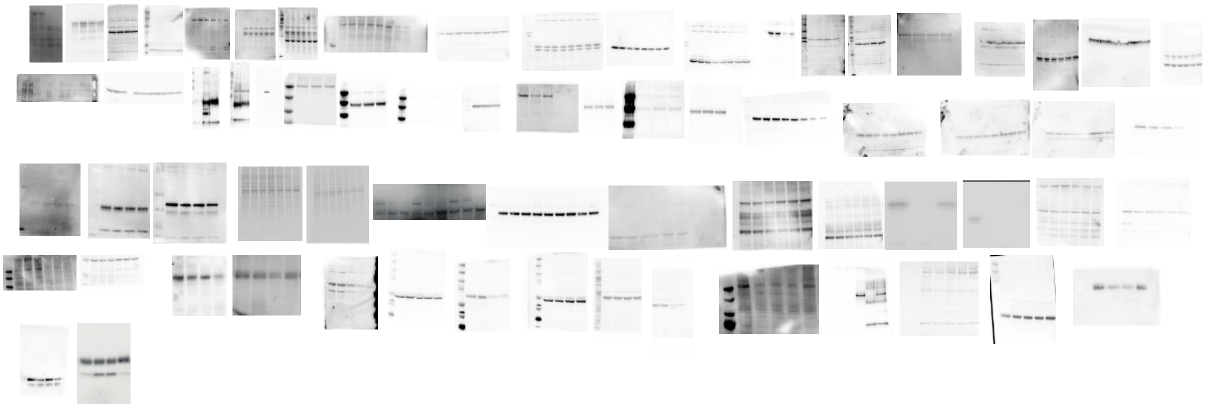

Supplementary Figures

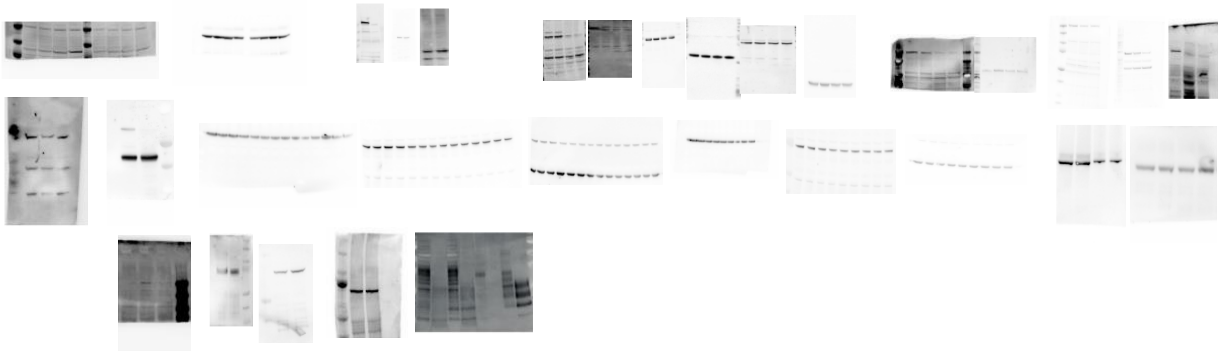

### Materials and Methods

*Mice and Intracerebral prion inoculations*

Mice were bred in high hygienic grade facilities and housed in groups of 3-5, under a 12 h light/12 h dark cycle (from 7 am to 7 pm) at 21±1°C, with sterilized food (Kliba No. 3431, Provimi Kliba, Kaiseraugst, Switzerland) and water *ad libitum*. Animal care and experimental protocols were in accordance with the Swiss Animal Protection Law, and approved by the Veterinary office of the Canton of Zurich (permits 130/2008 and 41/2012).

The following mice were used in the current study: C57BL/6, and *tga20* (B6;129-Tg(Prnp)a20Cwe Prnp<tm1Cwe>). Prions inoculations were performed by injecting different prion strains (RML6 and ME7) intracerebrally as described previously [3], with a dose corresponding to $3x10^5$ $LD_{50}$. Scrapie was diagnosed by the following set of clinical criteria: ataxia, kyphosis, priapism and hind leg paresis. Mice were sacrificed on the day of the onset of the clinical signs of scrapie. As control non-infectious brain homogenate (NBH) was intracerebrally injected into the mice and they were sacrificed approximately at the same time as when the prion infected mice reach the terminal stage (at the latest time point compatible with humane euthanasia) of the disease.

*Human Samples*

All human brain tissue samples used in the current study were anonymized and collected before 2010 and stored in a registered prion biobank at the University Hospital of Zurich. The study using the human brain tissue samples to monitor the levels of PIKfyve was approved by Kantonale Ethikkommision, Kanton Zürich (Permit ID: 2019-02431).

Brain lysates from post mortal CJD and control brain tissues were generated by slicing a small region of frontal cortex followed by homogenization in lysis buffer (10mM NaCl, 100mM Tris, 10mM EDTA, 0.5% sodium deoxycholate, 0.5% NP-40, pH adjusted to 7) using a TissueLyser LT. Total amount of protein was estimated using biciniconic acid (BCA) assay and 30µg of total protein was migrated on 4-12% SDS-PAGE and western blot was performed using anti-PIKfyve antibody.

*Cell culture and prion infection of cells*

Gt1 cells were grown in Dulbecco's modified eagle medium (DMEM) in the presence of 10% foetal bovine serum (FBS), penicillin-streptomycin and Glutamax (all obtained from Invitrogen). For prion infection of the cells, Gt1 cells growing in DMEM medium were incubated with either Rocky mountain laboratory strain of prion (RML6) prions (0.1%) or non-infectious brain homogenate (NBH; 0.1%) for 3 days in one well of a 6 well plate. This was followed by splitting the cells at 1:3 ratio every three days for at least 10 passages. The presence of infectivity in the cells was monitored by the presence of proteinase K (PK) resistant PrP, which can be detected by immunolabeling with POM1 antibody. At 70 dpi, the cells started developing vacuoles which were visualized by phase contrast microscopy. Transduction of lentivirus expressing GADD34 ($1 \times 10^8$ TU) was performed on chornically prion infected cells at 75dpi. Cells were lysed on 79dpi and processed for western blotting.

*Quantification of vacuoles*

Number of vacuolated cells were manually annotated and quantified using phase contrast microscopy. Four fields of vision were randomly chosen for each cover slip and 250 cells were counted per field of vision. In total 1000 cells were counted per experimental condition and the total number of vacuolated cells are represented in the figures. A Chi-Square test was carried out to estimate the statistical significance.

*Acyl Rac Assay*

In the first step of the Acyl rac methodology post nuclear supernatants were generated from the brains of terminally sick prion infected mice. Brains were homogenized in homogenization buffer (BH) (20 mM Hepes, pH 7.4, 200 mM NaCl, 1 mM dithiothreitol (DTT), 0.1 mM EDTA and 0.3 mM PMSF) by using a Teflon glass homogenizer. The homogenate was subjected to centrifugation at 5000rpm for 15 min. The supernatant obtained was transferred into a separate Eppendorf. The amount of protein in the sample was estimated using a BCA assay and 1 mg of the protein was subjected to ultracentrifugation at 450000rpm for 1 hour. After centrifugation, the supernatant was discarded and pellet was resuspended in 100µl of Buffer 1 (Hepes 25mM, NaCl 25mM, EDTA 1mM, pH 7.4) in the presence of 0.5% Triton X 100 and 1x protease inhibitors and vortex the sample. A further 200µl of blocking buffer (Hepes 100mM, EDTA 1mM, SDS 2.5%) was added to the sample

along with 1.5% of MMTS and incubated at 40°C for 4 hours with intermittent vortexing. 900µl of ice-cold acetone to the sample in the next step and incubated at -20°C for 20 minutes. Centrifuge the samples at 7500rpm for 10 min and decant the supernatant. Wash the pellet with 70% acetone (5 times) followed by centrifugation at 7500rpm for 10 min. Air dry the pellet and then re-dissolve it in 400µl of binding buffer (Hepes 100mM, EDTA 1mM, SDS 1.5%). Take 1/10$^{th}$ of the sample (40µl) as input. 150µl of the sample is treated with freshly prepared hydroxylamine and 150µl without any treatment. Incubate both tubes with 50µl of the activated beads and leave the sample overnight on the rotating wheel at 4°C. Wash the beads with binding buffer (3 times) followed by resuspension in the sample buffer (50µl) with 10mM DTT. The samples were loaded on the gels and western blots were performed using anti PIKfyve antibody. For controls antibodies against TRAPα and Calnexin were used.
To identify the DHHC enzyme responsible for acylating PIKfyve, Gt1 cells were transfected with siRNA against different DHHC (all from Qiagen) for 72hrs followed by cell lysis to obtain post nuclear supernatant. Once the post nuclear supernatant was generated the samples were treated in the exact way as described for the brain lysates.

*Western blot analysis*
Mice brains were homogenized using TissueLyser LT for 5 min in 10 vol of lysis buffer (0.5% Nonidet P-40, 0.5% 3-[(3-cholamidopropyl)dimethylammonio]-1-propanesulfonate (CHAPS), protease inhibitors (complete Mini, Roche), phosphates inhibitors (PhosphoSTOP, Roche) in PBS, and centrifuged at 1000g for 5 min at 4°C to remove debris. For Gt1 cells, lysates were prepared by exposing the cells to lysis buffer (PBS+Triton 1%+ Protease inhibitors) for 15 min at 4°C followed by scraping off the cells from the plates. The lysates were centrifuged at 8000rpm for 10 min and supernatant was used for further analysis. Protein lysates extracted from brains or cells were subjected to the standard BCA Assay (Thermofisher) to estimate the total amount of protein present and in all cases equal amounts of protein was loaded onto the SDS-PAGE (Novex NuPAGE 4-12% Bis-Tris Gels). For PIKfyve, p-TFEB, TRPML1 western blots, 80µg of total brain / cell lysate was loaded on the gels. For all the other proteins, 40µg of total protein was loaded onto the gels. Samples were transferred onto nitrocellulose membrane using iBlot (Invitrogen) according to the manufacturer's instructions. For proteins less than 100kDa the transfer was performed at 20V for 7 min. For proteins larger than 100kDa, the transfer was performed at 15V for 15 min. All samples were blocked in 5% SureBloc (lubio biosciences) for 1h followed by incubation with the primary antibody overnight in Tris-buffered saline-Tween (TBS-T) at 4°C. Secondary Peroxidase-Goat Anti-Mouse IgG (H+L) (#62-6520) or Peroxidase-Goat Anti-Rabbit IgG (H+L) (#111.035.045), used at 1:10000 1h at RT and membranes were developed using Luminata Crescendo (Millipore) and images were acquired using Fusion Solo 7S Edge (Vilber).

*Cerebellar organotypic cultures slices preparation*
Cerebellar organotypic cultured slices (COCS) were generated from 9-12 day old pups with a thickness of 350µm as described previously [4]. Cultures were maintained in a standard cell incubator (37 °C, 5% $CO_2$, 95% humidity) and the culture medium was changed three times per week. For prion inoculation of COCS, freshly generated slices were treated with 100µg of prion infected brain homogenate for every 10 slices followed by culturing in a 6-well Millicell-CM Biopore PTFE membrane insert (Millipore) according to previously published protocol[5]. Infected slice cultures were maintained for a maximum of 45 days when neurodegeneration is prominent in the slices. For treatment with GSK2606414, slices were incubated with 20µM of the drug for 14 days with a change of medium and replenishment with a fresh stock of drug every 3 days. On Day 35, the COCS were subjected to lysis and protein and RNA were isolated for further experiments.
For prion mimetics, toxicity in slices was induced by exposure to toxic anti- $PrP^C$ antibodies targeting the globular domain, such as single chain POM1 antibody, after a 14-day recovery period, according to previously published protocol [6]. *tga20* COCS were exposed to POM1 (67 nM), or as a control with IgG (67nM) for 4 days and cell lysates for western blots were generated in the lysis buffer (PBS+Triton 1%+1x protease inhibitors).

*Immunofluorescence*
Gt1 cells were grown on coverslips (10mm) and on 8 well culture slides (BD Biosciences) for prion infected cells, washed with ice cold 1X PBS and fixed with 4% paraformaldehyde (PFA) in PBS, pH

7.4, for 15 min at RT. After washing 2X with ice-cold PBS, cells were permeabilized with 0.1% Triton X-100 in PBS for 10 min at RT followed by rinsing in 1xPBS for three times and incubated for 1 hour in blocking solution (10% FBS in PBS) at room temperature. Cell were incubated in primary antibodies prepared in blocking buffer for 1 hour followed by washing with blocking buffer (3x) and further incubation with Alexa conjugated secondary antibodies (Invitrogen, Molecular Probes), prepared in the blocking buffer at a dilution of 1:5000 for 30 min along with 1 ug/ml DAPI (4',6-diamidino-2-phenylindole, Invitrogen). Samples were mounted onto the glass slides with fluorescent mounting medium (Dako). In the case of prion infected cells, glass slides were tightly sealed from the outside environment using nail polish. Samples were imaged using a CLSM Leica SP5 ZMB. Images were processed and analyzed using the "Image J" software.

*Transmission electron microscopy*
Transmission electron microscopy was performed as previously described[7]. COCS samples were fixed *in situ* with 2.5% glutaraldehyde + 2% paraformaldehyde in 0.1 M phosphate buffer, pH 7.4 and embedded in Epon. Ultrathin sections were mounted on copper grids coated with Formvar membrane and contrasted with uranyl acetate/lead citrate. Micrographs were acquired using a Hitachi H-7650 electron microscope (Hitachi High-Tech, Japan) operating at 80 kV. Brightness and contrast were adjusted using Photoshop.

*Flow Cytometry*
Gt1 cells were plated in 6 well plates at a dilution of 250,000 cells/well and were transfected with either a control siRNA or siRNA targeting PIKfvye the following days. Cells were left in the medium containing siRNA for up to 72 hours and the generation of the vacuoles was visualized using phase contrast microscopy. Cells were washed 1x with FACS buffer (PBS +10% FBS) followed by incubation with PBS-EDTA (2 mM) for 15 min for detachment. Once the cells were detached, pooled quadruplicates were resuspended in FACS buffer and subjected to centrifugation at 900rpm for 5 min. The pellets were resuspended in FACS buffer and the samples proceeded to flow cytometry. Before sample acquisition, each sample was treated with 5 µM LysoSensor Yellow/Blue (LysoSensor™ Yellow/Blue DND-160, ThermoFisher, #L7545) for 4.5 min at 37°C according to the manufacturer's guidelines. Consecutively, 5 µl of 1:5000 pre-diluted SYTOX Red dead cell stain (SYTOX™ Red dead cell stain, ThermoFisher, #S34859) was added to each sample, shortly vortexed and further incubated for 30 s followed by immediate sample acquisition and recording. Acquisition was performed using a BD FACSAria™ Fusion. Optical configurations were set as follows. A 355 nm UV and a 633 nm Red laser were used for optimal excitation of LysoSensor Yellow/Blue and SYTOX Red dead cell stain, respectively. The emission of LysoSensor Yellow/Blue was recorded using a LP502 mirror in combination with a BP530/30 filter and a LP410 mirror in combination with a BP450/20 filter. The emission of SYTOX Red was recorded using a BP670/30 filter. The flow cytometry data was analyzed using FlowJo 10.6.1. Gt1 cells were first gated for singlets and for living cells (SYTOX Red negatives) followed by further depiction in BP530/30 (acid) and BP450/20 (neutral/basic) histograms. The ratio of the mean fluorescence intensity (MFI) 530/30 of LysoSensor-stained minus MFI 530/30 LysoSensor-unstained and MFI 450/20 of LysoSensor-stained minus MFI 450/20 LysoSensor-unstained cells was calculated and plotted for the time-points 24, 48 and 72 hours post transfection (Figure 4B).

*Pulse-chase assays*
To monitor the half-life of PIKfyve, Gt1 cells were treated with thapsigargin or DMSO as a control, for 3 hours followed by incubation in the starvation medium (DMEM without methionine and cysteine) for 40 min along with thapsigargin to deplete the endogenous stores of methionine and cysteine. Cells were then labeled with 50µCi/ml $^{35}$S-methionine/cysteine for 20 min followed by a chase in normal medium at different time points. After the chase, the cells were harvested an isotonic HEPES buffer (pH-6.8) containing 2%CHAPS and protease inhibitor cocktail. Post nuclear supernatants were obtained by centrifuging the sample at 10000g for 10 min. PIKfyve was immunoprecipitated with anti PIKfyve antibody (Sigma) followed by incubation with Protein G dynabeads (Invitrogen) for 2h at 4°C. The immunoprecipitates were migrated on a 4-12% Tris-BIS gels followed by fixation and drying of the gels. The dried gels were exposed to phosphoscreen and the radiolabeled products were revealed using a Fuji film.

*Immunoprecipitation*

Protein extraction from prion infected or C57BL/6 mice brains was performed using mechanical lysis in IP buffer (HBS buffer (pH 6.8) with 2% CHAPS and cocktail of protease inhibitors (Roche)). Once lysed, the samples were subjected to centrifugation at 10000g for 10 min. Protein content in the sample was estimated using BCA assay (with BSA as a standard) and 1mg of the protein was used for the immunoprecipitation assays. For the cell lines, lysis was performed for 20 min at 4°C in IP buffer (1% Triton X-100 in PBS, Protease inhibitors 1x) followed by centrifugation for 10 min at 10000g. In both cases supernatants were precleared and incubated for 16 h at 4°C with antibodies and Protein G dynabeads (Invitrogen). After immunoprecipitation, the beads were washed for three times with the IP buffer and resuspended in the sample buffer (2x) after the final wash. The samples were heated at 95°C for 5 min and migrated on 4-12% Tris-Bis gels with the MOPS buffer.

*RNAi and shRNA*

Gt1 cells were plated at a density of 500,000 cells/well of a 6 well plate and 24h later were washed with PBS and replenished with medium without antibiotics. siRNA (final conc of 25 pmol/ well) was transfected into the cells using RNAi Max reagent (1%) according to manufacturer's instructions. 72 hours later cells processed to either obtain RNA or protein lysates or fixed for staining followed by imaging. The following are the list of siRNA used in the current study:

| Gene name | siRNA Target sequence (5´-3´) |
|---|---|
| Zdhhc1 | CCTGGTCCTAAAGGGATTA |
| Zdhhc2 | GGCACCATTTGCCAATTGT |
| Zdhhc3 | GGGCATAGAACAATTGAAA |
| Zdhhc4 | GTCCGACCTAGAGAAATAT |
| Zdhhc5 | GCCCAAGATTGAAGACAAA |
| Zdhhc6 | CTGGGTGTTATAGCAATAT |
| Zdhhc7 | GGGTGTTTCAGGGAATCAT |
| Zdhhc8 | CCTGCTCTATGTGCTCAAT |
| Zdhhc9 | GGCTCTTGATAATGTTTGA |
| Zdhhc11 | GTGCACTTGATCGCAATTA |
| Zdhhc12 | GGGAGTTCATATCTTCACA |
| Zdhhc13 | GCTGGTAGAAGCAGGATAT |
| Zdhhc14 | GAGGCTGTAATATGCTTCT |
| Zdhhc15 | GCAGGTGTTTGGCGATAAT |
| Zdhhc16 | GGCCATTGCTTATCTGTGT |
| Zdhhc17 | GCGACACAATATGGAATAT |
| Zdhhc18 | GGCAGACAGTGAAACTCAA |
| Zdhhc19 | GGGTCCCAATTACATGTCT |
| Zdhhc20 | GCCCTTCCAAAGAGTTCTA |
| Zdhhc21 | CCACCAGGGTTTCTTTAAA |
| Zdhhc22 | GCCCTTCTCTTGTGTTGAT |
| Zdhhc23 | GCGGGTTACTTCTGATACT |
| Zdhhc24 | GTGTGGGCTTCCATAATTA |
| PIKfyve | AAGGGTGAACTAGACAATA |
| FIG4 | TTCGACATCTTTGAAGATG |

| | |
|---|---|
| VAC14 | CAGACTGAAGACTGTCTGA |
| TFEB | GGCAGAAGAAAGACAATCA |

shRNA targeting PIKfyve was generated by cloning the target sequence (5´-GGCTTATGTATGCTTGATG-3´) into pSuper.retro.puro plasmid between BglII and Xho1 sites. Gt1 cells were transfected with shRNA against PIKfyve followed by antibiotic selection (puromycin; 3µg/ml for 24h) of the transfected cells. At 144h post transfection cells were fixed and stained using anti TRPML1 antibody. For the vacuolation rescue experiment, cells were supplemented with bPIP (20µg/ml) at 120h post transfection with shRNA against PIKfyve for 24h followed by manual quantification of number of vacuolated cells.

*Hexosaminidase beta assay*

A fluorimetric assay was performed to detect the amount of Hexosaminidase beta in the samples using the manufacturer's instructions in Beta Hexosaminidase Activity Assay kit from Cellbiolabs. The assay was based on the principle that in the presence of hexosaminidase beta the substrate p-nitrophenol-N-acetyl-beta-D-glucosaminide is converted to p-Nitrophenol which can be measured at 450nm. 50µg of brain lysates from control mice and prion infected mice in a final volume of 50µl was incubated with 50µl of substrate solution at 37°C for 15 min followed by addition of neutralization solution (100µl). For cell lysates from prion infected and control Gt1 cells, 100µg of cell lysate was used.

*mRNA isolation and quantitative real time PCR*

Total RNA from brains of prion infected and controls were isolated using RNeasy Plus Universal Mini Kit (QIagen), according to the manufacturer's manual. After reverse transcription (QuantiTect Rev. Transcription Kit, QIagen), cDNA was processed for real time q-PCR using SYBR-green (Roche) and determination of ΔΔCT-values was done on a ViiA 7 real-time system (Applied Biosystems). Total RNA from Gt1 cells infected with prions was isolated using RNeasy Mini Kit (Qiagen), according to the manufacturer's manual. RNA levels of GAPDH were used to standardize expression levels. RT-PCR was performed using SYBR-green (Roche) and determination of ΔΔCT-values was done on a ViiA 7 real-time system (Applied Biosystems). For the primer sequences used in this study, see the table below:

| **Gene name** | **Forward primer (5´-3´)** | **Reverse primer (5´-3´)** |
|---|---|---|
| **PIKfyve** | AGTCTGTGAGGTCGCCTGTA | TCTGGAGTCTAACACTGAAGAGC |
| **Zdhhc1** | ATCCTTCTGGGCCTGCTTTC | CATCTCCTGAATGGACCGCA |
| **Zdhhc2** | TGCAGAGAAAGAATTGCTGGAG | CACAATATCGGATTGCGCCG |
| **Zdhhc3** | CCTCCAGATTGACGATGCCA | CACACTTCTCTGGCTGGAGG |
| **Zdhhc4** | CGGAAGTCTTGGGGAGCG | TCAGATAGCAGCTCCGCTTG |
| **Zdhhc5** | GCACCATATCCCATCTGCATC | TCCCTGTTTCCAGGTTAGTCA |
| **Zdhhc6** | CGACCTAGTCGACCCAGCATC | GAAACACAGCAGGACAAACGTC |
| **Zdhhc7** | AAGCAACTTGTAAGGGTGTTTC | CCACGTCATGACAGCACAGA |
| **Zdhhc8** | GCACACGCTGGTTAAGAAGG | GCAGATGAGTGGTGGTCAGT |
| **Zdhhc9** | GACGTGAGGAGCGTTCCATT | GTGGTAGCGACTTCTCCCTG |
| **Zdhhc11** | GTGGCATCAACAAGAACTGGG | AGGGCGTGGTAGGAACTTCT |
| **Zdhhc12** | CTGCACGACACCGAGCTAC | GGGGTCCATGAGTGACACAG |
| **Zdhhc13** | GATCAGTTGGGTGGTGACCT | CTGAAACAGGACCGCCAGAT |
| **Zdhhc14** | GCCTCCTGTGTGACAATGAC | GGCTGTTTTGAATTATTCCTTACCA |
| **Zdhhc15** | GATAGTCGAGATTATCCAGAAGGCT | TGGAACACCCGAACTGTGAC |
| **Zdhhc16** | GACTGCTCACTCAGCCTCTG | CTGCCCTTGGAAGCTCTTGA |
| **Zdhhc17** | CTGAAACGCTTTCTCCCAGC | CCCGTTTCGGTCTCGTACTC |
| **Zdhhc18** | TGCAGTCTGATACCGCGTTG | CAGGTGGCAAGCACATTCAG |
| **Zdhhc19** | GCGTGTTTGCTGCCTTCAAT | CCAGCCACCTACAAGGGAAT |
| **Zdhhc20** | TGGACGAATGAACCAACAGT | CAACCAGCCAGCAATGGTAG |

| | | |
|---|---|---|
| **Zdhhc21** | GGCATGCCGACACCCACT | TCGTATGGGCCTCCCTTCAAT |
| **Zdhhc22** | CCTCTACACCTCTCTGGCCT | CTCCGGAGAAGAACTGGCTG |
| **Zdhhc23** | CAACAACCGCACACTGAAGG | ACAATGATGGTCCATTCTCCGT |
| **Zdhhc24** | GCTTATCAGCTGCTCAATCTGC | GGTCACGACGTAGGATGCAC |
| **Cathepsin A** | TCAGGCAGTGAAAACTCGGG | CGGTTCCGGGCATGTCTTG |
| **Cathepsin B** | GCTCTTGTTGGGCATTTGGG | ACTCGGCCATTGGTGTGAAT |
| **Glucocerebrosidase A** | TGGAGAGAAGTGTGCTGGTG | CAGACCACTGAGCTGTAGCC |
| **LAMP1** | GCCCTGGAATTGCAGTTTGG | TGCTGAATGTGGGCACTAGG |
| **Galactosidase Alpha** | CCCGAGAGGGATTCAAAGGG | TGTGGACGTAATTTGCGAGGT |
| **Mucolipin 1** | TGCTGTGGACCAGTACCTGA | GTAGTACCGCTGGCAGAGAG |
| **GALNS** | CATGGACGATATGGGGTGGG | CTGCAGCCATCCGGTCTAAA |
| **Aryl Sulfatase B** | TGCGCCGATTGAGTCTTTGA | AACAGTGGTTTCTCCGGTGG |
| **ATP6V0E1** | GGGTCCTAACCGGGGAGTTA | ACAGAGGATTGAGCTGTGCC |
| **TPP1** | CTACTGGGTGGTCAGCAACA | CAGCCGTGGGTTACATCAAAG |
| **STAT3** | GCAATACCATTGACCTGCCG | ACGTGAGCGACTCAAACTGC |
| **βactin** | CTGAGCTGCGTTTTACACCC | CGCCTTCACCGTTCCAGTTT |
| **GAPDH** | CCACCCCAGCAAGGAGAC | GAAATTGTGAGGGAGATGCT |
| **PIKfvye variant1** | GCCACATCCTCAGGAGAG | GCGTTTCAATACTGTGCTG |
| **PIkfyve variant-2** | CTCCAGAAGGAAAGCAG | CAGTAGGTGCATGTCGG |
| **PIKfyve variant-3** | CATCCTCAGGAGAGCACAG | GAGGCGTTTCAATACTGTG |

*Antibodies*

The following primary antibodies were used in the present study.

| Antibody | Source |
| --- | --- |
| PIKfyve | Sigma |
| β Actin | Abcam |
| CaV1.2 | Alomone Labs |
| VAC14 | Proteintech |
| FIG4 | Sigma |
| LAMP1 | Abcam |
| LAMP2 | Abcam |
| PERK | Cell Signaling |
| (p)PERK | Cell Signaling |
| eIF2α | Cell Signaling |
| (p)eIF2α | Cell Signaling |
| TRPML1 | Alomone Lab |
| SARA | Abcam |
| GM130 | Abcam |
| HSP60 | Abcam |
| Calnexin | Enzo Life sciences |
| Calreticulin | Abcam |
| Flag | Sigma |
| TFEB | Bethyl laboratories |
| TFEB-pSer142 | Sigma |

*Secondary antibodies*
All HRP tagged secondary antibodies were obtained from Jackson laboratories and all Alexa tagged fluorescent secondary antibodies used for immunofluorescence and IHC were obtained from Invitrogen.

*Data availability*
All original data has been included in the article. Any additional information / data required will be made available by the corresponding author upon reasonable request.
